# Defective mechanosensory transduction of the new inner hair cells prevents hearing recover in the damaged cochlea

**DOI:** 10.1101/2023.02.18.529042

**Authors:** Xiang Li, Minhui Ren, Yunpeng Gu, Tong Zhu, Yu Zhang, Jie Li, Chao Li, Guangqin Wang, Lei Song, Zhenghong Bi, Zhiyong Liu

## Abstract

Hearing loss is a major health problem worldwide. Numerous attempts at regenerating functional hair cells (HCs) have been unsuccessful, but little is known about the main barrier that prevents us from achieving it and improving the hearing ability after damage. Here, we developed an *in vivo* genetic mouse model, by which the inner HCs (IHCs), the primary sound receptors innervated by the auditory neurons, were specifically damaged and the neighboring nonsensory supporting cells (SCs) were transformed into IHCs by ectopic expression of transient Atoh1 and permanent Tbx2. Despite ∼477 new IHCs were regenerated per cochlea and their differentiation status was more advanced than reported previously, no significant hearing improvement was achieved. By taking advantage of this unique model, we further found that the new IHCs expressed the functional marker vGlut3, harbored the similar transcriptomic profiles and electrophysiological properties as the endogenous IHCs. However, the mechanosensory transduction (MET) current could not be recorded in the new IHCs. Thus, our study indicated that the defective MET should be the main barrier that stops us from restoring the hearing capacity in the damaged cochlea and would pave the way for regenerating IHCs *in vivo*.

## INTRODUCTION

Sound detection is achieved through hair cells (HCs) in the cochlea, the auditory organ located in the ventral part of the inner ear. The cochlear auditory epithelium, also known as the organ of Corti (OC), harbors one row of inner HCs (IHCs) on its medial side and three rows of outer HCs (OHCs) on the lateral side (1–4). OHCs are sound amplifiers and specifically express Prestin, a motor protein encoded by *Slc26a5* (5, 6), and *Slc26a5^-/-^* mice display severe hearing impairment (7). Conversely, IHCs are primary sensory cells and specifically express vGlut3, which is encoded by *Slc17a8*, and *Slc17a8^-/-^* mice are completely deaf (8, 9) because vGlut3 is essential for the release of the excitatory neurotransmitter glutamate through the unique ribbon synapse structure between IHCs and auditory spiral ganglion neurons (SGNs) (10). Notably, both OHCs and IHCs are critical for sound detection and their damage causes permanent deafness in mammals, which have lost HC regeneration capacity (11–13). Intriguingly, the b-HLH transcription factor Atoh1 is necessary for cochlear pan-HC development: whereas all HCs disappear in *Atoh1^-/-^* mice (14–16), *Atoh1* overexpression induces supernumerary new HCs (17–21). Furthermore, Insm1 and Ikzf2 play crucial roles in specifying or stabilizing the OHC fate (22, 23), and our group and two other groups have independently shown that Tbx2 is required in IHC fate specification, differentiation, and maintenance (24–26).

Adjacent to HCs, the following distinct subtypes of supporting cells (SCs) are located from the medial to lateral side: inner border cells (IBCs), inner phalangeal cells (IPhs), pillar cells (PCs), and Deiters’ cells (DCs) (4). IBCs and IPhs belong to the medial SC subgroup and their gene-expression profiles and developmental origins are distinct from those of lateral PCs and DCs (27, 28). Whereas *Prox1* is transiently expressed and *Fgfr3* is permanently expressed in PCs and DCs (20, 27, 29–31), proteolipid protein 1 (encoded by *Plp1*) and solute carrier family 1 member 3 (*Slc1a3*) are enriched in IBCs and IPhs (32, 33). However, IBCs and IPhs currently cannot be readily distinguished based on a specific molecular marker, and thus we roughly grouped these cells as IBCs/IPhs in this study. PCs/DCs and OHCs have been suggested to be born from the same lateral progenitors (27), and, accordingly, we have previously shown that PCs and DCs can be converted into OHC-like cells through concurrent expression of Ikzf2 and Atoh1 (34). Similarly, IBCs/IPhs and IHCs share identical medial progenitors (27, 28), and we have reported that neonatal IBCs/IPhs are reprogrammed into IHCs expressing the early marker Myo6 but not vGlut3 through ectopic expression of Atoh1 alone (35), but into Myo7a+/vGlut3+ IHCs through dual expression of Tbx2 and Atoh1 (24). However, it remains unclear whether functional IHCs can be regenerated in the lesioned cochlea where endogenous IHCs are damaged, mimicking the HC repair that naturally occurs in nonmammalian vertebrates such as chicken and fish (11, 36). If not able to regenerate the functional IHCs and restore the hearing, what the main barrier is.

By exploiting a well-designed genetic mouse model, we first damaged endogenous neonatal IHCs and then conditionally induced Atoh1 and Tbx2 expression specifically in neonatal IBCs/IPhs. Approximately 477 new IHCs were yielded per cochlea. Both single-cell transcriptomic and electrophysiological assays showed that the new IHCs resemble the endogenous IHCs. However, the mechanosensory transduction is defective in the new IHCs, which should be one of the main barriers accounting for the failure to achieve significant hearing improvement in our current model. Our results highlight the future directions of how to regenerate function IHCs and realize hearing restoring *in vivo*.

## RESULTS

### Generating new *Fgf8*-DTR/+ model for IHC-specific damage

To achieve specific IHC damage in neonatal cochleae, we generated a new knockin mouse model, *Fgf8*-P2A-DTR-T2A-DTR/+ (briefly, *Fgf8*-DTR/+), in which diphtheria toxin (DT) receptor (DTR) expression is strictly controlled by *Fgf8* promoters and enhancers (Supplemental Figure 1A-C). *Fgf8* is highly and exclusively expressed in IHCs within perinatal cochleae (37–39), and transient DT administration would cause the death of DTR-expressing HCs (40–43). Notably, we inserted two DTR fragments to induce efficient IHC death by DT treatment at a dosage as low as possible, which theoretically would produce fewer side effects and allow better survival of mice as compared to higher dosages. Southern blotting confirmed the absence of random insertion of the targeting vector in the mouse genome (Supplemental Figure 1D and E), and tail-DNA PCR readily distinguished the wild-type (WT) and knockin alleles (Supplemental Figure 1F). Both heterozygous and homozygous mice were fertile and healthy and did not exhibit any abnormalities. These findings also indicated that Fgf8 expression itself was intact because *Fgf8^-/-^* mice die at ∼E9.5 (44). Thus, we predicted that IHCs and OHCs would be normal in *Fgf8*-DTR/+ mice in the absence of DT treatment (no DT), and that IHCs, but not OHCs, would be lost upon administration of DT (15 ng/g body weight) at P2.

Our predication of IHC loss was experimentally confirmed at P42 (Figure 1A-B’), and IHC quantification across entire cochleae revealed that only 16.3 ± 6.7 (n=4) IHCs were present in DT-treated *Fgf8*-DTR/+ mice at P42, significantly fewer than the 732.0 ± 15.4 (n=3) IHCs in no-DT littermates (n=3) (Figure 1C); thus, ∼97.8% of IHCs were lost following a single administration of DT at P2. Moreover, considerable IHC loss occurred in all cochlear turns (Figure 1D). Accordingly, auditory brainstem response (ABR) assays revealed that hearing thresholds at all frequencies in DT-treated mice (n=7) were drastically higher than those in no-DT mice at P42 (n=8) (Figure 1E). Notably, the three rows of OHCs were aligned well, as expected, in both no-DT and DT-treated *Fgf8*-DTR/+ mice (Figure 1A-B’). Collectively, our results indicated that *Fgf8*-DTR/+ is a powerful genetic model to specifically damage IHCs in the neonatal mouse cochlea.

**Figure 1.**
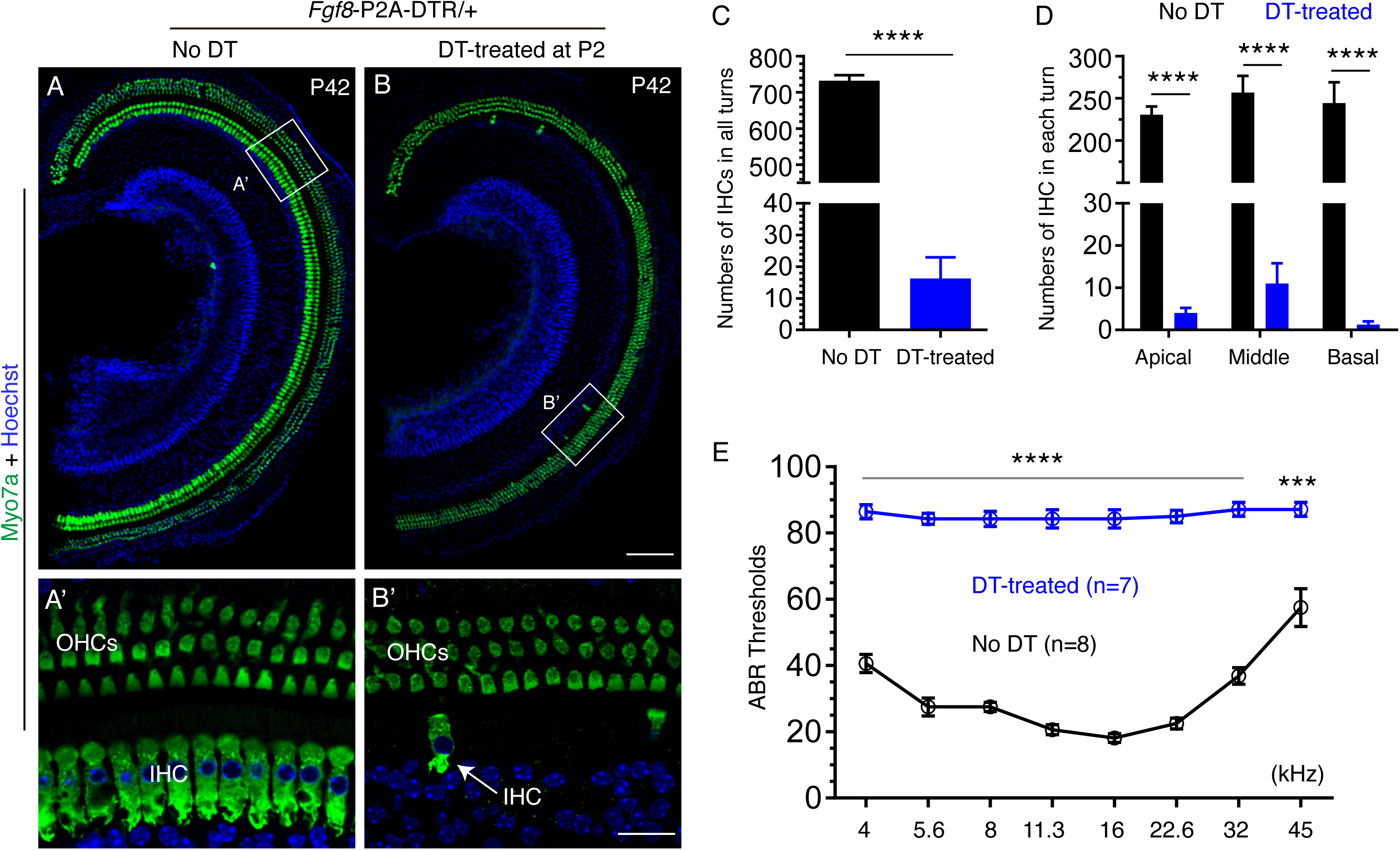
Specific damage of neonatal cochlear IHCs causes severe hearing impairment in *Fgf8*-P2A-DTR/+ mice. **(A-B’)** Staining of pan-HC marker Myo7a in cochleae of P42 *Fgf8*-P2A-DTR/+ mice that were either not treated with DT (no DT; A and A’) or treated with DT at P2 (DT-treated; B and B’). Boxed areas in (A) and (B) are shown at high resolution in (A’) and (B’), respectively. Almost all IHCs were lost after DT treatment (B’); arrow: one remaining IHC. Notably, OHCs were normal in both groups of mice. **(C)** Quantification of total IHC numbers in no-DT (black, n=3) and DT-treated (blue, n=4) *Fgf8*-P2A-DTR/+ mice at P42. Data are presented as means ± SEM. **** p<0.0001. **(D)** IHC quantification as in (C), except that data are presented for different cochlear turns. **(E)** ABR measurements compared between no-DT (black line) and DT-treated (blue line) *Fgf8*-P2A-DTR/+ mice. ABR thresholds at all frequencies in DT-treated mice were significantly higher (worse hearing) than those in no-DT mice. Data are presented as means ± SEM. *** p<0.001, **** p<0.0001. Scale bars: 200 μm (B) and 20 μm (B’).

### Damaging endogenous IHCs promotes reprogramming efficiency of Atoh1 and Tbx2

We previously showed that 29.5% ± 1.2% of neonatal cochlear IBCs/IPhs are converted into vGlut3-expressing IHCs in the undamaged cochlea upon dual expression of Atoh1 and Tbx2 (24), which is achieved using Plp1-CreER+; *Rosa26*-CAG-Loxp-Stop-Loxp-Tbx2*3×V5-P2A-DHFR-Atoh1*3×HA-DHFR-T2A-tdTomato*/+* (Plp1- TAT in short) (Supplemental Figure 1G). Plp1-CreER+ is a recognized IBC/IPh- specific Cre driver within the OC (33), and TAT denotes <u>T</u>bx2 (fused with V5 tag), <u>A</u>toh1 (fused with HA tag and protein unstable domain derived from dihydrofolate reductase (DHFR)), and tdTomato. If a protein of interest is fused with DHFR which contains the destabilizing domains (DDs), it will undergo rapid proteasomal degradation (45). However, trimethoprim (TMP), a cell-permeable small molecule, can bind to and stabilize the DDs, by which TMP treatment results in the protein degradation being stopped in a rapid, reversible, and TMP dose-dependent manner (45). Briefly, permanent Tbx2 and transient Atoh1 expression (hereafter Tbx2 and Atoh1 expression, in short) and permanent tdTomato expression are conditionally and specifically induced in neonatal IBCs/IPhs upon exposure to tamoxifen (TMX) at P0 and P1 and then TMP (for stabilizing Atoh1 transiently) at P3 and P4 (Supplemental Figure 1G). Thus, the tdTomato expression helps distinguishing new IHCs from the endogenous IHCs that do not express tdTomato. Notably, the success of transient Atoh1 expression is confirmed using the four lines of evidence described in our previous study (24), and a single dose of TMP can maximally stabilize Atoh1 protein for up to three days.

We determined here whether specific damage of endogenous IHCs further boosts the reprogramming efficiency of Atoh1 and Tbx2 (Figure 2A). Plp1-TAT; *Fgf8*-DTR/+ (Plp1-TAT-DTR, in short) mice were either (1) administered DT only and used as the control group (Figure 2B-B’’); or (2) administered TMX at P0 and P1, DT at P2, and TMP at P3 and P4, and defined as the TMX/DT/TMP-treated group (Figure 2C-C’’). Importantly, to accurately label IBCs/IPhs for subsequent fate-mapping analysis, TMX was administered before DT treatment (Figure 2A), because the opposite order could potentially alter the Cre expression pattern in Plp1-CreER+.

**Figure 2.**
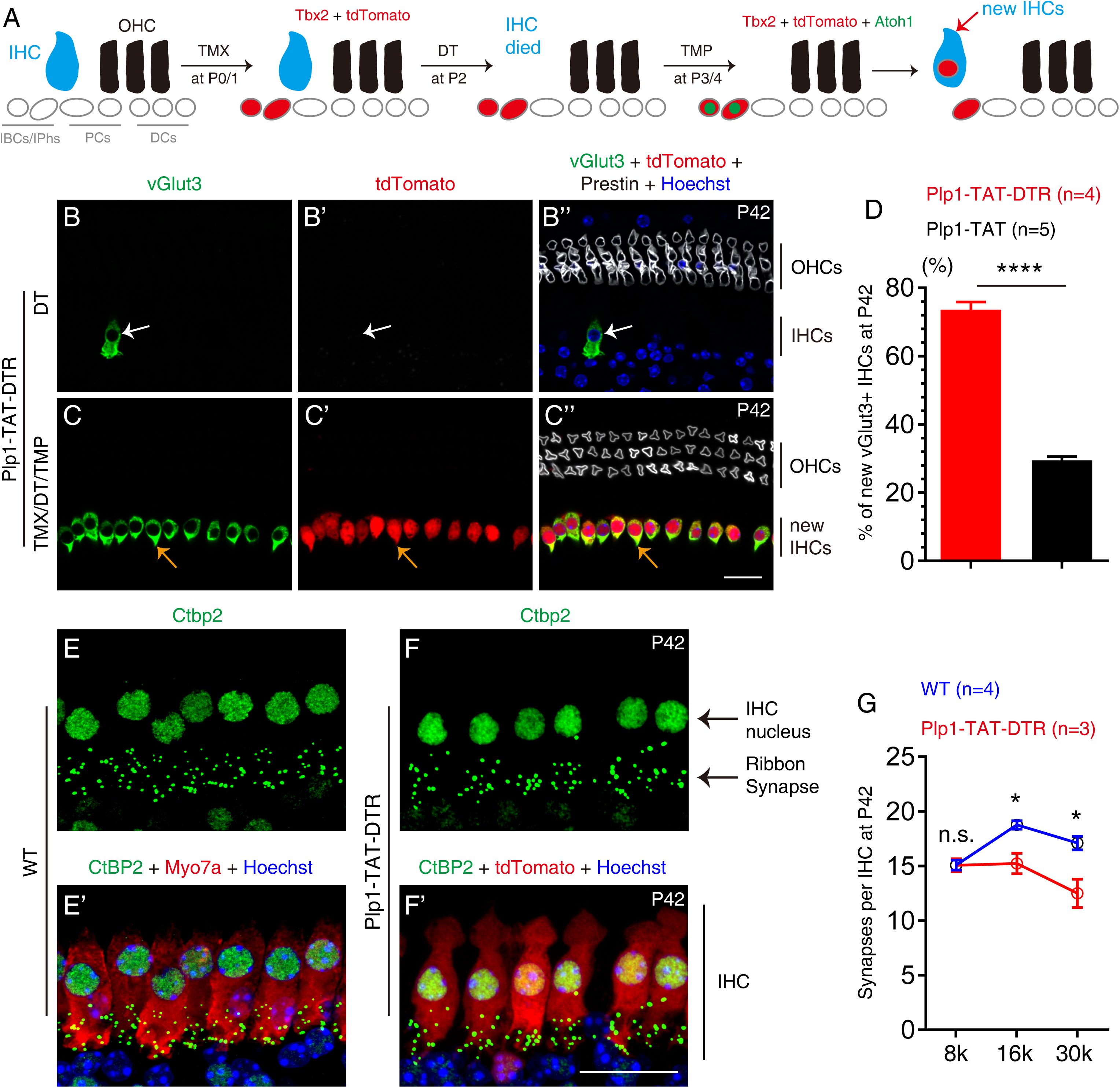
Damaging endogenous IHCs boosts reprogramming efficiency of Atoh1 and Tbx2. **(A)** Simple cartoon illustrating key cellular events during cell-fate conversion from neonatal IBCs/IPhs into new IHCs in the presence of endogenous IHC damage. **(B-C’’)** Triple labeling for vGlut3, tdTomato, and Prestin in DT-treated mice (B-B’’) and TMX/DT/TMP-treated mice (C-C’’) at P42. Arrows in (B-B’’): one remaining endogenous IHC that expressed vGlut3+ but not tdTomato and Prestin; arrows in (C-C’’): one new IHC that was tdTomato+/vGlut3+/Prestin-. Similar labeling is shown at lower magnification in Supplemental Figure 2A-B’’’. **(D)** Comparison of percentages of tdTomato+/vGlut3+ new IHCs between Plp1-TAT-DTR (red, n=4) and Plp1-TAT (black, n=5) mice at P42. Data are presented as means ± SEM. **** p<0.0001. Significantly higher percentages of tdTomato+ cells expressing Atoh1 and Tbx2 were converted into new IHCs in Plp1-TAT-DTR mice (with IHC damage) than in Plp1-TAT mice (without IHC damage). **(E-G)** Double staining of Ctbp2 and Myo7a in wild-type mice (WT; E-E’) and double labeling for Ctbp2 and tdTomato in Plp1- TAT-DTR mice (F-F’) at P42. (G) Synapse numbers were calculated by averaging numbers of Ctbp2+ puncta per Myo7a+ endogenous IHC (E-E’) or per tdTomato+ new IHC (F-F’). At 8, 16, and 30 kHz positions, we counted 70, 86, and 58 endogenous IHCs in WT mice (n=4) and 74, 79, and 57 new IHCs in Plp1-TAT-DTR mice (n=3), respectively. Synapse numbers were compared between endogenous and new IHCs at three cochlear positions. Data are presented as means ± SEM. * p<0.05. New IHCs harbored fewer synapses than endogenous IHCs at 16 and 30 kHz. Scale bar: 20 μm (C’’ and F’).

As in DT-treated *Fgf8-DTR/+* mice (Figure 1B and B’), only 17.3 ± 4.0 (n=4) IHCs were present in DT-treated Plp1-TAT-DTR mice at P42 (arrows in Figure 2B-B” and Supplemental Figure 2A-A’’’). By contrast, 477.0 ± 48.0 (n=4) new IHCs, which were tdTomato+/vGlut3+/Prestin- (Figure 2C-C’’ and Supplemental Figure 2B-B’’’), were detected in entire cochleae of TMX/DT/TMP-treated Plp1-TAT-DTR mice at P42. Intriguingly, the number of vGlut3+ new IHCs here (477.0 ± 48.0) was ∼2.2-fold higher than the corresponding number (221.8 ± 19.1) in Plp1-TAT mice in which endogenous IHCs were not damaged (24). Accordingly, normalizing the new IHC numbers against the total number of tdTomato+ cells (close to new IHCs) revealed that 73.5% ± 2.3% of the tdTomato+ cells (which were originally IBCs/IPhs) were reprogrammed into vGlut3+ new IHCs in TMX/DT/TMP-treated Plp1-TAT-DTR mice, which was also markedly higher than the 29.5% ± 1.2% measured in Plp1-TAT mice (Figure 2D). Based on these results, we concluded that the reprogramming efficiency of Atoh1 and Tbx2 in neonatal IBCs/IPhs was drastically increased when endogenous IHCs were damaged.

### Ribbon synapses form between new IHCs and SGNs

Next, we determined whether ribbon synapses were formed between new IHCs and SGNs. Ctbp2+ ribbon synapses were quantified based on co-staining for Ctbp2 and Myo7a in WT mice (Figure 2E and E’) and co-labeling for Ctbp2 and tdTomato (Figure 2F and F’) in TMX/DT/TMP-treated Plp1-TAT-DTR mice at P42. The average numbers of synapses per new IHC were similar (at 8 kHz) or slightly lower (at 16 and 30 kHz) relative to those per endogenous IHC (Figure 2G).

Besides Ctbp2 staining, we performed triple labeling for neurofilament (NF)-200, tdTomato, and Myo7a, which revealed that the NF-200 signal was lower in DT-treated Plp1-TAT-DTR mice (Supplemental Figure 2C-C’’) than in TMX/DT/TMP-treated Plp1-TAT-DTR mice (Supplemental Figure 2D-D’’) at P42. However, the NF-200 signal was the strongest in the control Plp1CreER+; *Rosa26*-LSL-tdTomato (Ai9)/+ (Plp1-Ai9) mice that were also administered TMX and TMP (Supplemental Figure 2E- E’’). Furthermore, in DT-treated Plp1-TAT-DTR mice, the NF-200 signal appeared weaker in the regions where IHCs were completely lost (yellow arrows in Supplemental Figure 2C-C’’) than in regions near the remaining IHCs (red arrows in Supplemental Figure 2C-C’’). Collectively, our results supported the conclusion that ribbon synapses were formed between IHCs and SGNs in Plp1-TAT-DTR mice at P42, although the synapse numbers were lower than in the control Plp1-Ai9 mice.

### Vast majority of new IHCs are produced by direct transdifferentiation

In nonmammalian vertebrates such as chicken and fish, hearing ability can be restored after damage (12, 13, 36), and this occurs through two processes: one, transdifferentiation, where SCs located near damaged HCs directly switch their SC fate to HC fate without proliferation; and two, mitotic regeneration, where the SCs first proliferate and expand their populations before starting to transdifferentiate into auditory HCs. To ascertain which process is used in our model, we analyzed Plp1-TAT- DTR mice that, besides being treated with TMX at P0 and P1, DT at P2, and TMP at P3 and P4, were administered 5-ethynyl-2ʹ-deoxyuridine (EdU) either at P5 and P7 and analyzed at P9 (Supplemental Figure 3A-B’’’) or at P7, P9, and P11 and analyzed at P12 (Supplemental Figure 3C-D’’’).

In both experimental paradigms, no EdU incorporation was detected in the tdTomato+/vGlut3+ new IHCs (arrows in Supplemental Figure 3B-B’’’ and D-D’’’). Notably, however, successful EdU administration was confirmed by the presence of EdU+ cells in the cochlear SGN area (Supplemental Figure 3A and C). These results suggested that the vast majority of the new IHCs were derived from direct transdifferentiation rather than mitotic regeneration. Accordingly, limited cell proliferation was also recently reported in the process of Atoh1-mediated conversion of greater epithelial ridge (GER) cells into HCs (21).

### Ultrastructural analysis of new IHCs by using scanning and transmission electron microscopy

We first used scanning electron microscopy (SEM) to determine whether hair bundles or stereocilia were present on the surface of new IHCs at P42 in two genetic models: (1) Plp1-Ai9 (Figure 3A and A’) and (2) Plp1-TAT-DTR (Figure 3B-C’). In control Plp1-Ai9 mice, the typical stereocilia of endogenous OHCs exhibited a “V” or “W” shape (pink), and the stereocilia of IHCs displayed a “bird-wing” pattern (purple) (Figure 3A and A’). As expected, IHC stereocilia were absent in DT-treated Plp1-TAT- DTR mice at P42 (Figure 3B and B’), whereas the “bird-wing”-like stereocilia were recovered in TMX/DT/TMP-treated Plp1-TAT-DTR mice (Figure 3C and C’). Although the IHC “bird-wing”-like stereocilia in TMX/DT/TMP-treated Plp1-TAT-DTR mice were not as regularly organized as the stereocilia of endogenous IHCs, these stereocilia of new IHCs were generally better developed than those of the OHC-like cells described in our previous report (34). Nonetheless, this “bird-wing”-like morphology agreed with the IHC fate of the cells.

**Figure 3.**
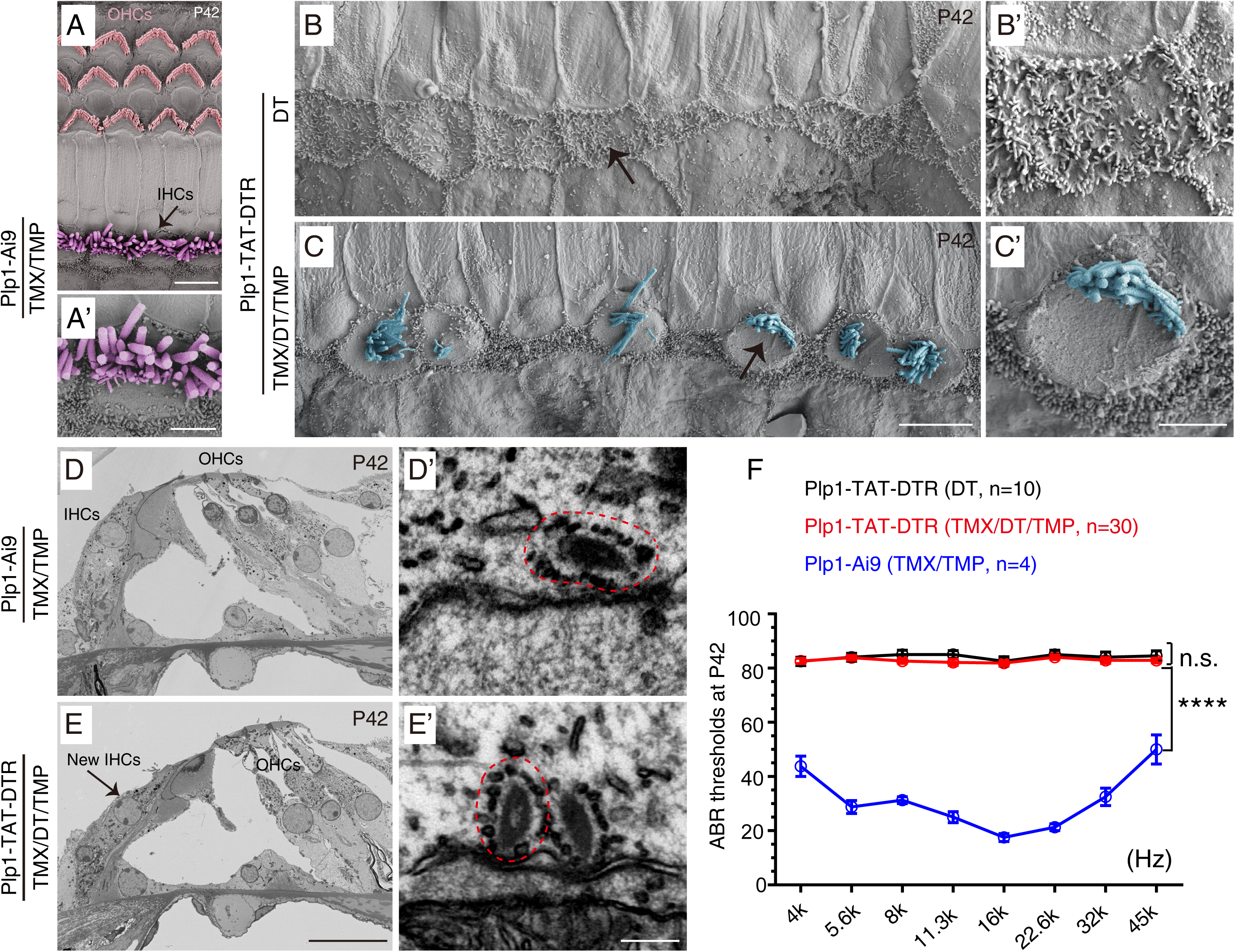
Stereocilia and ribbon synapses are present in new IHCs. (A-C’) SEM analysis of mouse models at P42: control Plp1-Ai9 (A-A’) and Plp1-TAT-DTR (B-C’). Plp1-TAT-DTR mice were further divided into two subgroups: treatment with DT only (DT, B-B’) and treatment with TMX, DT, and TMP (TMX/DT/TMP, C-C’). Arrows in (A, B, and C): areas shown in higher magnification in (A’, B’, and C’), respectively. As compared with stereocilia in endogenous IHCs (purple, A-A’), those in new IHCs (light blue, C-C’) were irregularly organized. **(D-E’)** TEM analysis of control Plp1-Ai9 mice (D-D’) and TMX/DT/TMP-treated Plp1-TAT-DTR mice (E-E’). Dotted red circles in (D’ and E’): ribbon synapses with high electron density. **(F)** ABR measurements of control Plp1-Ai9 mice (blue line) and DT-treated (black line) and TMX/DT/TMP- treated (red line) Plp1-TAT-DTR mice at P42. Data are presented as means ± SEM. **** p<0.0001. Scale bars: 5 μm (A, C), 2 μm (A’, C’), 20 μm (E), and 200 nm (E’).

We next used transmission electron microscopy (TEM) to further confirm the presence of ribbon synapses in new IHCs (Figure 3D-E’). Consistent with our immunostaining results (Figure 2 and Supplemental Figure 2), ribbon synapses featuring high electron density were captured in both control Plp1-Ai9 mice and TMX/DT/TMP-treated Plp1-TAT-DTR mice (red dotted circles in Figure 3D’ and E’). However, the auditory brainstem response (ABR) measurements did not reveal significant hearing improvement in TMX/DT/TMP-treated Plp1-TAT-DTR mice (n=30), relative to that in DT-treated Plp1-TAT-DTR mice (n=10) (Figure 3F). Notably, the hearing thresholds at all frequencies in the 30 Plp1-TAT-DTR mice were also significantly higher than those in control Plp1-Ai9 mice (blue line in Figure 3F) at P42. Thus, we concluded that in the 477.0 ± 48.0 new IHCs, further manipulations are necessary to achieve improved hearing recovery.

### New IHCs exhibit similar electrophysiological features with endogenous IHCs, but they harbor the defective mechanosensory transduction

We performed whole-cell patch-clamp recording of individual new IHCs or WT IHCs to verify cellular functions specific to IHCs at the single-cell level. Both endogenous IHCs in WT mice and new IHCs in Plp1-TAT-DTR mice were recorded at P42. First, both WT IHCs (Figure 4A) and new IHCs (Figure 4B) exhibited voltage-dependent Ca^2+^ currents, although current-amplitude variations were larger in the new IHCs (n=10, cell number) than in WT IHCs (n=5) (Figure 4C). Second, linear capacitance, which is indicative of cell size, of new IHCs (n=9) was smaller than that of WT IHCs (n=9) (Figure 4D), which is consistent with the new IHCs being derived from IBCs/IPhs, both originally smaller than endogenous IHCs. Third, exocytosis- triggered change in membrane capacitance, a readout for the release of glutamate-containing vesicles, was measured when depolarization was induced in both WT and new IHCs (Figure 4E), and, moreover, surface-area increments in WT and new IHCs were comparable (Figure 4F), which indicated that normal sustained vesicle-release functions were established in the new IHCs.

**Figure 4.**
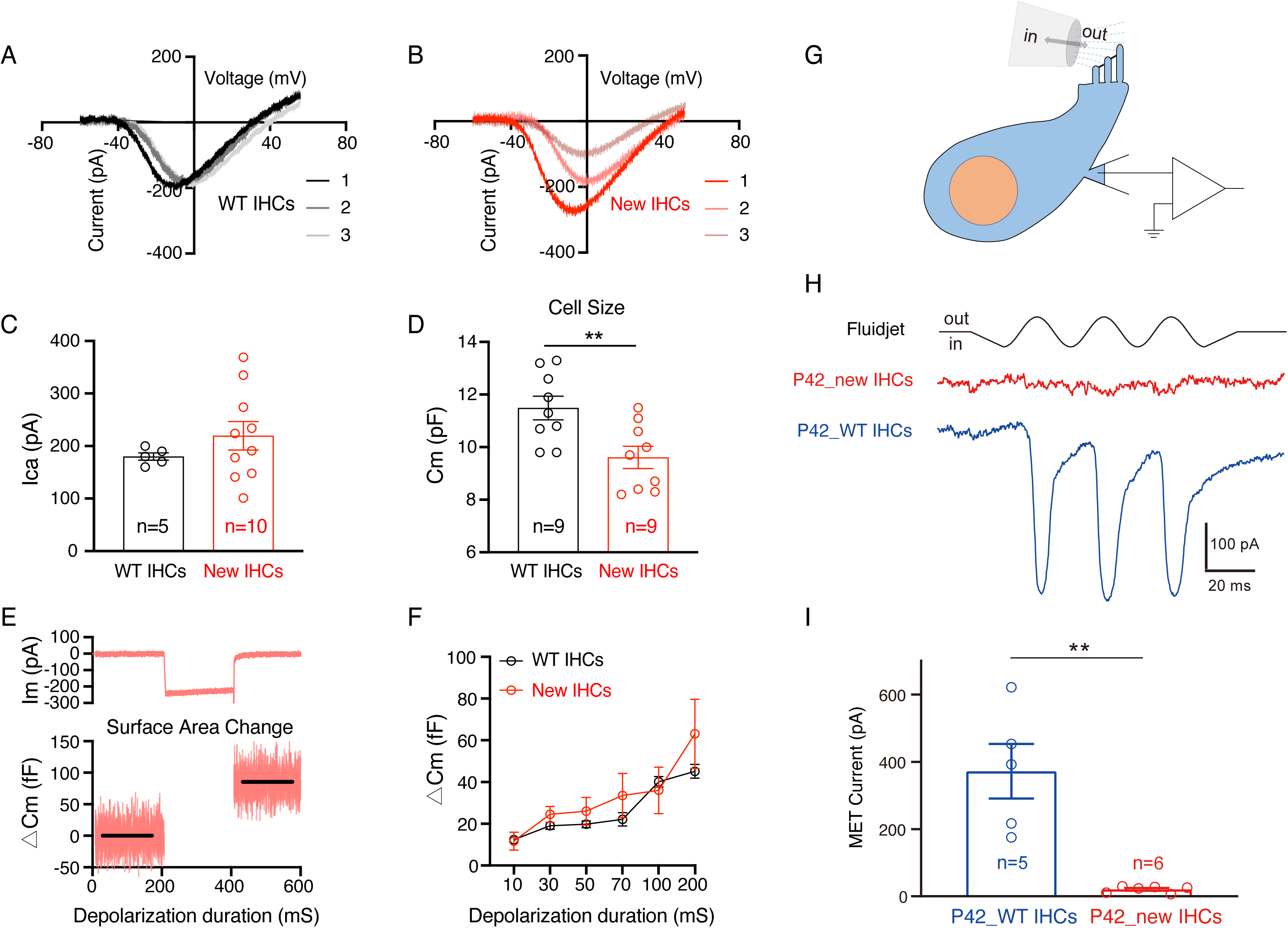
New and wild-type endogenous IHCs exhibit similar electrophysiological characteristics. (A-C) Calcium current–voltage relationships measured in wild-type (WT) IHCs (A) and new IHCs (B). Three WT IHCs (A) and three new IHCs (B) were chosen to present, respectively. Quantification and statistical analysis of individual maximal Ca^2+^ current amplitudes are compared in (C). No significant difference was detected. Notably, current variations were larger in new IHCs than in WT IHCs. Three current–voltage curves out of 5 measured for WT IHCs and 10 for new IHCs are plotted in (A) and (B), respectively. **(D)** Comparison of linear capacitance (cell size) measurements of 9 WT IHCs and 9 new IHCs; new IHCs were significantly smaller than WT IHCs. Data are presented as means ± SEM. ** p<0.01. **(E)** Representative current (upper panel) and C_m_ (lower panel) change triggered by a 200 ms depolarizing stimulus from -80 to 0 mV voltage step. ΔCm: IHC surface-area increment due to exocytosis. **(F)** Exocytic ΔCm (surface-area change) elicited by depolarizing stimuli of 10–200 ms duration in WT (black) and new (red) IHCs. **(G)** Simple cartoon illustrating how fluidjet-evoked MET current is recorded in IHCs. **(H)** Representative MET currents in new IHCs (red) and wild-type (WT) IHCs (blue) at P42. **(I)** Quantification of MET currents in new IHCs and WT IHCs. Data are presented as means ± SEM. ** p<0.01.

We also measured mechanosensory transduction (MET) currents in the new IHCs (Figure 4G). At P42, IHCs (n=5, cell number) from WT mice exhibited fluidjet- dependent MET currents (blue color in Figure 4H and I), whereas only very weak (low signal-to-noise ratio) or background MET currents were induced in the new IHCs (n=6) (red color in Figure 4H and I). It should account for the failure to achieve significant hearing improvement in the Plp1-TAT-DTR mice (Figure 3F). Nonetheless, because new IHCs exhibited voltage-dependent Ca^2+^ currents and neurotransmitter release, we hypothesized that the global gene-expression profiles of the new IHCs would diverge from those of both neonatal IBCs/IPhs (origin of new IHCs) and adult IBCs/IPhs (final fate of neonatal IBCs/IPhs if ectopic Atoh1 and Tbx2 are not expressed). Conversely, the new IHCs could resemble adult endogenous IHCs. To test this, we performed single-cell RNA-seq analysis.

### Global gene-expression profiles of neonatal and adult IBCs/IPhs are distinct

Initially, we first obtained the transcriptomic profiles of adult endogenous IBCs/IPhs at P42 (P42_IBCs/IPhs, in short). Cochlear tissues of Plp1-Ai9 mice at P42 were subject to 10× Genomics single-cell RNA-seq. A total of 4,051 qualified cells were obtained and divided into 18 clusters by using t-distributed stochastic neighbor embedding (t-SNE) analysis (Supplemental Figure 4A), and an average of 3,133 genes were detected per cell; tdTomato was enriched in the cells of Clusters 0, 7, 10, and 12, confirming that these were cells of the Plp1+ lineage (Supplemental Figure 4B), and the cells of Clusters 0, 7, and 12 were cochlear glial cells. Here, we only focused on Cluster 10 cells, in which *Epcam* and *Slc1a3* were enriched (Supplemental Figure 4C and D): Epcam is a marker specifically expressed in the cochlear epithelium (46–48), and *Slc1a3* encodes a glutamate-aspartate transporter (Glast), a known marker of IBCs/IPhs (32, 35). Ultimately, we selected 55 cells in Cluster 10 and defined these as P42_IBCs/IPhs. Second, we identified 110 neonatal endogenous IBCs/IPhs at P1 (P1_IBCs/IPhs, in short) from one previous single-cell RNA-seq study (27). The detailed criteria used for defining P1_IBCs/IPhs and P42_IBCs/IPhs are described in the methods section.

Next, we mixed the P1_ and P42_IBCs/IPhs and performed uniform manifold approximation and projection (UMAP) analysis (Supplemental Figure 4E and F); two separate clusters were formed, suggesting distinct transcriptomic profiles of P1_and P42_IBCs/IPhs. Furthermore, trajectory analysis revealed the developmental track of IBCs/IPhs between P1 and P42 (Supplemental Figure 4G and H); the presence of two distinct subclusters at each age supported the notion that the IBCs/IPhs show intrinsic differences.

### Identification of 432 IHC- and 423 IBC/IPh-enriched genes

We next aimed to identify IHC- or IBC/IPh-enriched or specific genes. Because adult cochlear HCs cannot be readily captured using the droplet-based microfluidics platform (27, 49), smart-seq-based transcriptomic profiles of 50 endogenous mature IHCs at P30 (P30_IHCs in short) were obtained by manually picking tdTomato+ IHCs from the *vGlut3*-P2A-iCreER/+; *Rosa26*-LSL-tdTomato (Ai9)/+ mice reported in our previous study (24). Here, our analysis included the P30_IHCs and P1_IBCs/IPhs and P42_IBCs/IPhs, as well as 16 cochlear PCs and DCs at P60 (P60_PCs/DCs) from another study (34). UMAP analysis revealed that the majority of the P30_IHCs aggregated together (green in Supplemental Figure 5A). Furthermore, the P42_IBCs/IPhs and P60_PCs/DCs were divergent, indicating that the global gene-expression profiles of adult IBCs/IPhs (OC medial side) and PCs/DCs (OC lateral side) were also highly different from each other. Altogether, the results highlighted the high quality of our single-cell RNA-seq data.

We sought to identify genes significantly enriched in P42_IBCs/IPhs (“IBC/IPh genes”) and P30_IHCs (“IHC genes”) and use these genes to assess the degree of cell-fate conversion in vGlut3+ new IHCs. However, transcriptomic profiles obtained from 10× Genomics and smart-seq approaches cannot be directly compared. Thus, IBC/IPh and IHC genes were defined as follows. As in our previous study (34), our criteria for regarding a gene of interest as being expressed at a “high” level in the smart-seq approach were an average transcripts per million (TPM) value of >16 and expression by at least half the cells of the gene at TPM > 16. First, we selected 3,349 genes that were highly expressed in P30_IHCs (green circle in Supplemental Figure 5B). Second, we selected 3,373 genes (black circle in Supplemental Figure 5B) that were highly expressed in adult IHCs by reanalyzing data from one previous bulk RNA-seq study (50). Notably, 2,226 genes overlapped between the 3,349 and 3,373 genes from the two datasets and were regarded as genes that can be considered with confidence to be highly expressed in adult IHCs (Supplemental Figure 5B and purple circle in Supplemental Figure 5C). Third, 4,060 genes were highly expressed in PCs and DCs (black circle in Supplemental Figure 5C), and 1,781 genes overlapped between the 4,060 and 2,226 genes; these were genes shared by adult PCs/DCs and IHCs. By subtracting these 1,781 genes from the 2,226 genes, we identified 445 genes expressed in P30_IHCs but not P60_PCs/DCs (orange lines in Supplemental Figure 5C and orange circle in Supplemental Figure 5D). Fourth, we performed transcriptomic comparison between the 55 P42_IBCs/IPhs and all other non-IBC/IPh populations (Supplemental Figure 4), which revealed that 436 genes were significantly enriched (log_2_FC>0.5, p<0.05) in P42_IBCs/IPhs (black circle in Supplemental Figure 5D).

Intriguingly, 13 genes including *Tbx2* were shared by IHCs and IBCs/IPhs, which agrees with Tbx2 also being expressed in postnatal IBCs/IPhs (24). By further excluding these 13 genes, we ultimately defined 432 IHC genes (red lines in Supplemental Figure 5D) and 423 IBC/IPh genes (blue lines in Supplemental Figure 5D). Notably, these IHC genes included both IHC-specific genes such as *Slc17a8*, *Otof*, and *Slc7a14* and pan-HC genes such as *Myo7a*, *Tmc1*, and *Rbm24* (green arrows in Supplemental Figure 5E and F). Similarly, *Slc1a3* and other pan-SC genes such as *Sox2*, *Sox10*, and *Hes1* were included among the IBC/IPh genes (blue arrows in Supplemental Figure 5E and F). Supplemental Table 1 presents the entire list of the 432 IHC and 423 IBC/IPh genes, as well as the 13 genes shared by P30_IHCs and P42_IBCs/IPhs.

### Transcriptomic profiles of new IHCs and differentiating endogenous IHCs

Next, we determined the transcriptomic profiles of new IHCs by applying 10× Genomics single-cell RNA-seq to cochlear samples from TMX/DT/TMP-treated Plp1- TAT-DTR mice at P42. A total of 4,661 qualified cells were obtained and further divided into 19 clusters (Supplemental Figure 6A), and an average of 2,862 genes were detected per cell. Briefly, the 41 cells in Cluster 15, which were tdTomato+/Myo7a+/vGlut3+ (arrows in Supplemental Figure 6B-D), were defined as new IHCs (P42_new IHCs, in short). Moreover, tdTomato expression in the P42_new IHCs confirmed that the cells were derived from Plp1+ neonatal IBCs/IPhs. As in Plp1-Ai9 mice (Supplemental Figure 4), very few adult endogenous IHCs (Myo7a+/Slc17a8+/tdTomato-) were captured (Supplemental Figure 6A-D). Our success in capturing P42_new IHCs, but not adult endogenous IHCs, suggested that the new IHCs better tolerate the droplet-based microfluidics platform.

We also obtained single-cell transcriptomic profiles of endogenous IHCs at E16 (E16_IHCs), P1 (P1_IHCs), and P7 (P7_IHCs) from one recent single-cell RNA-seq study (27). These E16_IHCs (n=106 cells), P1_IHCs (n=103), and P7_IHCs (n=79) were annotated in the trajectory line (Supplemental Figure 6E and F). Altogether, we have collected transcriptomic profiles of the three key cell types at multiple ages: endogenous IBCs/IPhs (P1 and P42), endogenous IHCs (E16, P1, P7, and P30), and new IHCs (P42).

### New IHCs drastically upregulate IHC genes and downregulate IBC/IPh genes

We next directly calculated the key molecular-signature differences between P42_new IHCs and P42_IBCs/IPhs, because both datasets were generated using the 10× Genomics single-cell RNA-seq platform. Expression levels of 626 genes were significantly higher (log_2_FC>0.5 and p<0.05) in P42_new IHCs than in P42_IBCs/IPhs (Supplemental Figure 7A), and gene ontology (GO) analysis revealed that these genes were enriched in functions involved in, for example, sensory perception of sound, auditory receptor cell differentiation, and vesicle-mediated transport (Supplemental Figure 7B); the genes enriched in each GO category are included in Supplemental Table 2. Furthermore, 186/626 (29.7%) genes were among the 432 IHC genes and included the IHC-specific genes *Slc17a8*, *Otof*, and *Slc7a14* and the pan-HC genes *Cib2*, *Lhfpl5*, *Espn*, *Myo6*, and *Myo7a* (pink arrows in Supplemental Figure 7A). All 626 and 186 genes are listed in Supplemental Table 3. Thus, we concluded that 43.1% (186/432) of the IHC genes were substantially upregulated in P42_IHCs. Conversely, the expression levels of 337 genes were significantly lower (log_2_FC>0.5 and p<0.05) in P42_new IHCs than in P42_IBCs/IPhs (Supplemental Figure 7A), and GO analysis revealed that these genes were overrepresented in positive regulation of cell proliferation and negative regulation of the Notch pathway (Supplemental Figure 7C); the genes in each GO category are included in Supplemental Table 2. More importantly, 191/337 (56.7%) genes were included among the 423 IBC/IPh genes, such as *Slc1a3*, *Gjb2*, *S100a1*, *Sox2*, and *Sox10* (blue arrows in Supplemental Figure 7A); these 337 and 191 genes are all listed in Supplemental Table 3. Therefore, we concluded that 45.2% (191/423) of the IBC/IPh genes, which otherwise would be expressed in P42_IBCs/IPhs, were significantly repressed in P42_new IHCs.

### Global gene-expression profile of new IHCs resembles that of adult endogenous IHCs

To determine the differentiation status of P42_new IHCs, we analyzed a mixture of the P42_new IHCs (n=41) with E16_IHCs (n=106), P1_IHCs (n=103), P7_IHCs (n=79), and P30_IHCs (n=50): Six cell clusters were formed, with 98% (49/50) of the P30_IHCs belonging to Cluster 3 and 100% (41/41) of the new P42_new IHCs sorting into Cluster 4 (Figure 5A), and, notably, the P42_new IHCs and P30_IHCs aggregated close to each other (dotted circle in Figure 5B). Furthermore, Clusters 0 and 1 primarily included E16_IHCs and P7_IHCs, respectively, and P1_IHCs sorted into Clusters 2 and 5, which might reflect the heterogeneous differentiation status of IHCs in the basal-apical gradient. Thus, we concluded that the global gene-expression profile of P42_new IHCs most closely resembles that of P30_IHCs, even though these cells were assigned to two distinct clusters. Moreover, trajectory analysis of all IHCs together revealed the maturation process from E16_IHCs to P30_IHCs (black arrows in Figure 5C), and the calculated pseudotime matched the genuine ages of the IHC lineage (Figure 5D).

**Figure 5.**
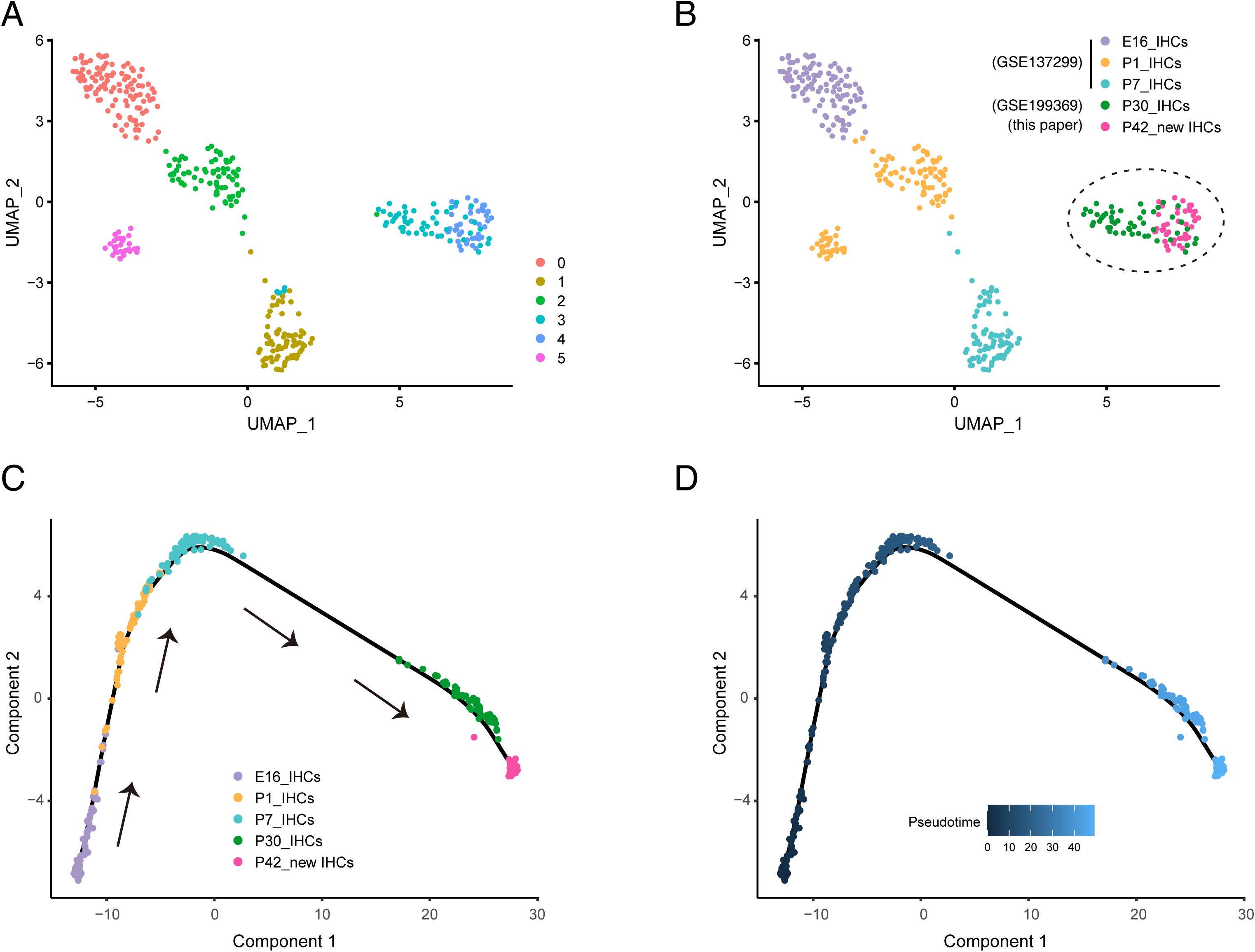
New IHCs resemble adult endogenous IHCs in transcriptomic analysis. (A-B) UMAP analysis of cell mixture including E16_IHCs, P1_IHCs, P7_IHCs, P30_IHCs, and P42_new IHCs; 6 main cell clusters formed (A), and P42_new IHCs and P30_IHCs clustered close to each other (dotted circle in B). **(C-D)** Trajectory analysis and pseudotime analysis of 5 cell populations listed in (B). Black arrows: calculated differentiation track of endogenous IHC lineage. P42_new IHCs and P30_IHCs were again close to each other. Moreover, the calculated ages of the IHC lineage matched their genuine ages.

Collectively, our results supported the conclusion that P42_new IHCs resemble endogenous mature P30_IHCs the most.

### New IHCs minimally express utricle and OHC genes

Considering the similarity of P42_new IHCs and P30_IHCs, we predicted that utricle HC and OHC genes would be minimally expressed in these cells, which was confirmed by our analysis here. Besides the 26 neonatal utricle HC genes and 53 neonatal OHC genes that were reported in our previous study (34), we also identified 151 adult OHC-specific genes (24). The entire list of these 151 adult OHC genes, which included previously known OHC markers such as *Ocm*, *Slc26a5*, and *Ikzf2*, is presented in Supplemental Table 4. Briefly, we regarded a given utricle HC gene or OHC gene as being expressed in P42_new IHCs if these two criteria were both met: (1) the gene’s average post-normalized expression value was >0.5; and (2) >50% of the P42_new IHCs expressed the gene with an expression value of >0.5. However, the standard of expression level of >0.5 (10× Genomics platform) cannot be directly compared with the TPM value of >16 (smart-seq platform) described above.

According to the two criteria, we found that 15/151 (9.9%) adult OHC genes (Supplemental Figure 8A), 6/53 (11.3%) neonatal OHC genes (Supplemental Figure 8B), and 2/26 (7.7%) neonatal utricle HC genes (Supplemental Figure 8C) were expressed in P42_new IHCs. The detailed expression levels of the OHC genes or utricle HC genes in each P42_new IHC are included in Supplemental Table 5. Altogether, our results revealed that the P42_new IHCs minimally expressed utricle and OHC genes, in agreement with the cells most closely resembling mature P30_IHCs.

### Molecular difference between new IHCs and mature endogenous IHCs

Lastly, we characterized the disparity between P42_new IHCs and P30_IHCs. Because the gene-expression profiles of the cells were obtained using two distinct methods (10× Genomics and smart-seq), the profiles cannot be compared directly. Thus, we instead focused on the 432 IHC genes and 423 IBC/IPh genes (Supplemental Figure 5E and F and Supplemental Table 1) and assessed whether a given IHC or IBC/IPh gene was expressed in P42_new IHCs by adopting the same standard (normalized expression value of >0.5) as that used to determine expression patterns of the vestibular HC or OHC genes described above (Supplemental Figure 8).

In P42_new IHCs, 249/432 (57.6%) IHC genes were highly expressed and included IHC-specific genes such as *Otof*, *Slc17a8*, and *Slc7a14* and pan-HC genes such as *Lhfpl5*, *Lmo7*, *Pou4f3*, and *Myo7a* (green arrows in top half of Supplemental Figure 9A); conversely, the remaining 183/432 (42.4%) genes were defined as being expressed at low levels in P42_new IHCs (bottom half of Supplemental Figure 9A). All 249 and 183 genes are listed in Supplemental Table 6.

We hypothesized that if P42_new IHCs do not fully resemble P30_IHCs, the new IHCs would also maintain expression of a fraction of IBC/IPh genes. Accordingly, among the 423 IBC/IPh genes, 168 (39.7%) were maintained at a high expression level, including *S100a1*, *Id1*, and *Id4* (green arrows in Supplemental Figure 9B); by contrast, the remaining 255/423 (60.3%) genes were expressed at undetectable or low levels in P42_new IHCs and included genes such as *Sox10* and *Hes1* (gray arrows in Supplemental Figure 9B). These 168 and 255 genes are also listed in Supplemental Table 6.

In Figure 6, we summarize the upregulation and downregulation of IHC genes and IBC/IPh genes during cell-fate conversion, as well as the molecular difference between P42_new IHCs and adult endogenous IHCs. Currently, the overall transcriptomic similarity between P42_new IHCs and P30_IHCs is ∼60%. Future efforts aimed at diminishing the disparities between these cells are required to optimize the regenerated new IHCs.

**Figure 6.**
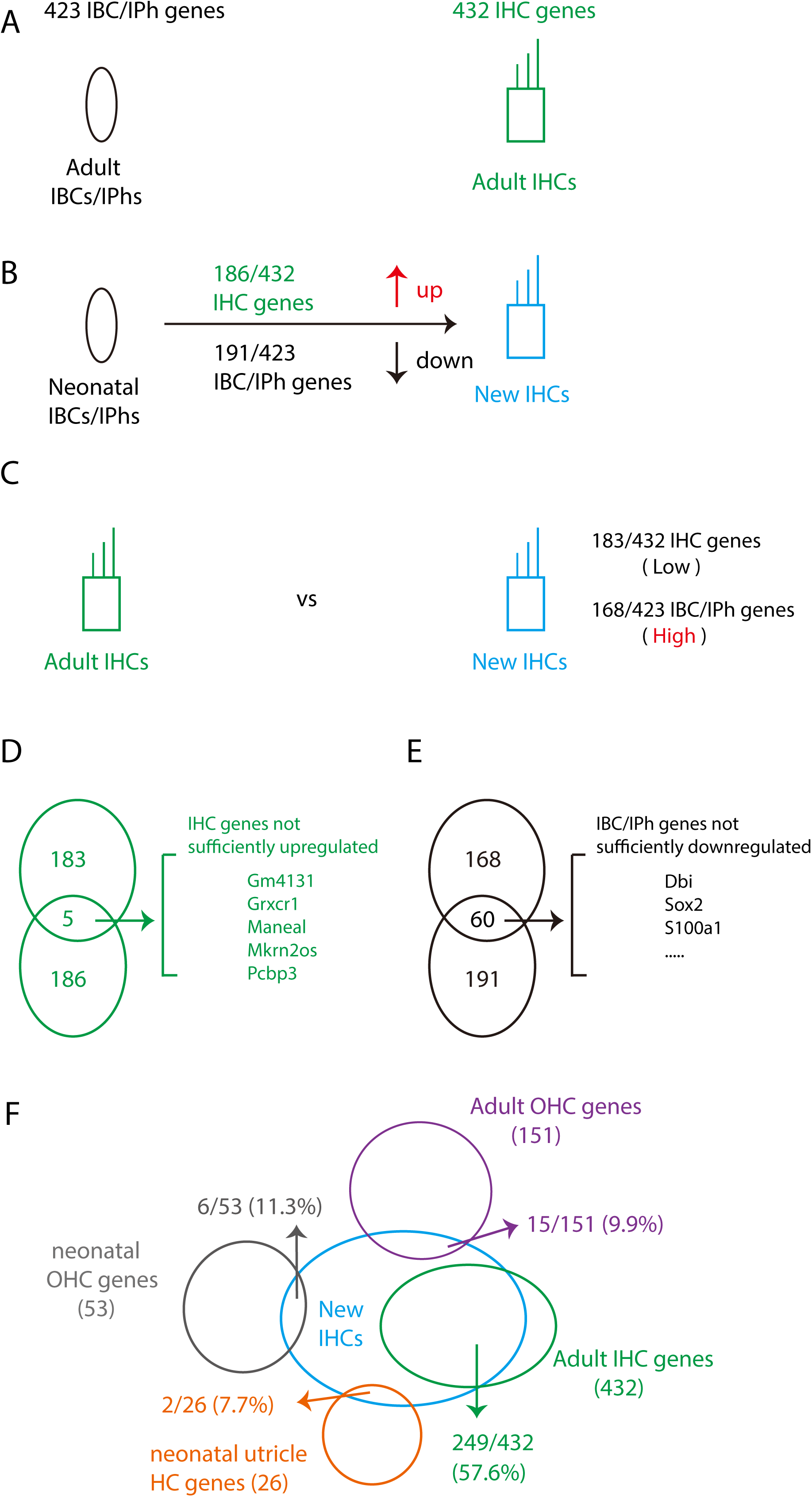
Simple summary of cell-fate conversion from neonatal IBCs/IPhs into IHCs. **(A)** A total of 423 IBC/IPh and 432 IHC genes were defined. **(B)** During cell- fate conversion from neonatal IBCs/IPhs into IHCs, 186 IHC genes were upregulated and 191 IBC/IPh genes were downregulated. **(C)** Relative to levels in adult endogenous IHCs, 183 IHC genes were expressed at significantly lower levels and 168 IBC/IPh genes were maintained at higher levels in new IHCs. **(D-E)** Illustration of 5 IHC genes that were not sufficiently upregulated in new IHCs (D) and 60 IBC/IPh genes that were not sufficiently downregulated in new IHCs (E). **(F)** Global similarity between new IHCs and adult endogenous IHCs was 57.6%. Accordingly, new IHCs minimally expressed neonatal and adult OHC or utricle HC genes.

## DISCUSSION

This study provided compelling evidence that in the presence of endogenous IHC damage, persistent Tbx2 coupled with transient Atoh1 expression efficiently transformed neonatal cochlear IBCs/IPhs into IHCs expressing the functional marker vGlut3. Notably, the new IHCs resemble the endogenous IHCs in molecular, ultrastructural, and electrophysiological aspects, except that the MET of the new IHCs is defective, which is the main barrier toward hearing improvement.

### *Fgf8*-DTR/+: a mouse model to damage endogenous IHCs

*Fgf8*-DTR/+ mice survive well after a single DT treatment even though Fgf8 is also expressed in other organs besides the ear. Our results clearly showed that the *Fgf8*- DTR/+ knockin strain is well suited for *in vivo* IHC damage in neonatal cochleae. In terms of damaging neonatal IHCs, *Fgf8*-DTR/+ offers advantages over the *vGlut3*- DTR/+ strain (51), because Fgf8 shows IHC-specific expression whereas vGlut3 is expressed in both IHCs and glial cells in neonatal cochleae (10, 39). However, *vGlut3*- DTR/+ is a valuable model for damaging adult IHCs, because *Fgf8* expression is turned off but vGlut3 expression is maintained in adult IHCs (10, 23). Including our previous *Slc26a5*-DTR/+ model designed to specifically damage postnatal cochlear OHCs (34), these three knockin mouse strains are sufficient to allow damage of cochlear IHCs or OHCs at all postnatal ages, depending on the experimental goals. Moreover, considering the expression pattern of Fgf8 in the embryonic cochlea (37, 38), we propose that *Fgf8-DTR/+* will likely serve as a favorable model for ablating IHCs soon after their birth and investigating how the loss of nascent IHCs affects overall cochlear development.

### Damaging endogenous IHCs potently augments efficiency of Atoh1 and Tbx2 in reprogramming neonatal IBCs/IPhs

Reprogramming of SCs into HCs is an effective process for hearing recovery after trauma in nonmammalian vertebrates (52). Previous work has shown that, relative to Atoh1 expressed alone (35), Tbx2 and Atoh1 together convert neonatal mouse IBCs/IPhs into IHCs not only more effectively, but also yield new IHCs in a more differentiated state (24). Although the molecular mechanisms underlying this phenomenon remain unclear, we speculate that Atoh1 and Tbx2 interact synergistically in cellular reprogramming, a notion supported by the finding that Atoh1, Gfi1, and Pou4f3 also convert cochlear GER cells or medial SCs into IHC-like cells more effectively than Atoh1 alone (21, 53).

Whether damaging endogenous IHCs can augment the reprogramming efficiency of Tbx2 and Atoh1 has remained unknown, and, to our knowledge, this is the first study to address this question. By using the new *Fgf8*-DTR/+ strain, we obtained results clearly showing that the reprogramming efficiency (73.5% ± 2.3%) of Tbx2 and Atoh1 expression in the presence of IHC damage was markedly higher than that in the absence of HC damage (29.5% ± 1.2%), and, as expected, larger numbers of new IHCs were generated in the presence of IHC damage than in its absence. How might IHC damage enhance the reprogramming efficiency of Tbx2 and Atoh1? We speculate that, one, certain unknown components released from the dying IHCs might stimulate the response of IBCs/IPhs to Tbx2 and Atoh1 manipulation; or two, intercellular signaling, such as Notch signaling (21), from IHCs to IBCs/IPhs might prevent the IBCs/IPhs expressing Atoh1 and Tbx2 from adopting the IHC fate, with these inhibitory effects being lost upon IHC ablation and yielding larger numbers of regenerated IHCs. Accordingly, ablating adult OHCs notably increased the ability of Atoh1 and Ikzf2 to reprogram PCs and DCs into OHC-like cells (34).

### Degree of cell-fate conversion from IBCs/IPhs into IHCs

Marked hearing improvement at all frequency regions was not achieved using our current experimental paradigm, which raises the question as to what extent the new IHCs resemble endogenous IHCs. We grossly addressed this question by quantifying the upregulation and downregulation of IHC genes and IBC/IPh genes, respectively, in the new IHCs (Figure 6A and B): 43.1% (186/432) of the IHC genes were significantly upregulated and 45.2% (191/423) of the IBC/IPh genes were significantly downregulated in P42_new IHCs, relative to P42_IBCs/IPhs, which suggested a cell-fate conversion of ∼45%; however, in the P42_new IHCs, 42.4% (183/432) of the IHC genes were still expressed at low levels, and, conversely, 39.7% (168/423) of the IBC/IPh genes were expressed at high levels (Figure 6C), suggesting a cell-fate conversion of ∼60%. Considering both calculations, we conclude that the cell-fate conversion rate lies between 45% and 60%. Notably, this estimation of cell-fate conversion is simplified and solely based on transcriptomic analysis.

Intriguingly, 5 genes overlapped between the 183 and 186 IHC genes mentioned above (Figure 6D) suggested that these 5 genes were not sufficiently upregulated to match their expression level in mature endogenous IHCs; conversely, 60 genes overlapped between the 168 and 191 IBC/IPh genes (Figure 6E), indicating that these 60 genes were not sufficiently downregulated. A similar phenomenon has been recorded in the cell-fate conversion from adult PCs/DCs into Prestin+ OHC-like cells (34). Together, these findings suggest that further upregulating IHC genes and downregulating IBC/IPh genes in new IHCs would facilitate the differentiation of the new IHCs.

### Proposed future approaches to promote more complete differentiation of new IHCs

Cochlear IHCs perform two main functions in the hearing process: signal detection through MET channels in stereocilia at their top surface (54–56), and signal transduction mediated by ribbon synapses at their bottom (57–59). Signal detection and transduction are both indispensable for normal hearing (60). Ribbon synapses were formed here between new IHCs and SGN fibers (Figure 2) and vesicle release was observed when the new IHCs were depolarized (Figure 4), which suggested normal signal transduction at the bottom part of the new IHCs. By contrast, two lines of evidence indicate that defective MET in the new IHCs represents one key barrier toward achieving marked hearing improvement in our current genetic model: (1) stereocilia on the apical surface of P42_new IHCs were irregularly aligned (Figure 3), and (2) the MET current in P42_new IHCs was very weak (Figure 4G-I).

The weak MET current in new IHCs (at least in the apical turn) noted above is not likely due to a lack of expression of MET components: key proteins involved in MET function, including Tmc1 and Lhfpl5 (61–63), are expressed in the new IHCs (Supplemental Table 3), much as the case in OHC-like cells in our previous study (34). It is not a surprise because MET-related proteins are expressed independently of the formation of stereocilia (64). Conversely, stereocilia were not as regularly organized in new IHCs as in endogenous IHCs (Figure 3C and C’), and thus the MET impairment is more likely caused by defective MET channel assembly. *Emx2* is reported to be required in hair bundle organization (65). Recently, the intrinsic spontaneous calcium actions potential activity is important in maintaining the integrity of stereociliary bundles of IHCs (66). However, methods to further optimize MET channel organization in the new IHCs currently remain unknown.

In future studies, we will identify key candidate genes that regulate the morphogenesis of IHC stereocilia by using our *in vivo* high-throughput genetic- screening approach (67, 68). If mutation of any these candidate genes results in the formation of stereocilia resembling those in P42_new IHCs, we will include the genes to the combination of *Atoh1* and *Tbx2*. Thus, combined ectopic expression of key IHC developmental genes in IBCs/IPhs could serve as a promising future approach to regenerate IHCs in patients with hearing impairment.

## MATERIALS AND METHODS

### Generation of *Fgf8*-DTR/+ knockin mouse strain

*Fgf8*-DTR/+ mice were created by co-injecting one sgRNA against the Fgf8 locus, donor DNA, and Cas9 mRNA into one-cell-stage mouse zygotes. The sgRNA was *5ʹ- AGCTGGGCGAGCGCCTATCG-3ʹ,* and the donor DNA was designed as described in Supplemental Figure 1. F0 mice born from pseudopregnant mothers were screened, and mice with potentially successful gene targeting were identified using junction PCR. All potentially correct F0 mice were crossed with WT C57BL/6 mice, and the born F1 mice that were germ-line stable were subject to further junction-PCR and Southern blotting analyses to confirm the absence of random insertion of donor DNA in the F1 genome. Southern blotting was performed according to our protocol described previously (10). F2 and subsequent mice were genotyped using regular tail-DNA PCR. The detailed primer sequences are included and described in Supplemental Table 7.

### Mouse breeding and drug treatment

Tamoxifen (TMX; Cat#: T5648, Sigma-Aldrich) was dissolved in corn oil (Cat#: C8267, Sigma-Aldrich). Diphtheria toxin (DT; Cat#: D0564, Sigma-Aldrich) was dissolved in 0.9% NaCl solution. Trimethoprim lactate salt (TMP; Cat#: T0667, Sigma-Aldrich) was resolved in 1× phosphate-buffered saline (PBS). In all experiments, mice were administered TMX (3 mg/40 g body weight) at P0 and P1, DT (15 ng/g body weight) at P2, and TMP (300 μg/g body weight) at P3 and P4. EdU labelling kit (Cat# C10637, Thermo Scientific) was used, following the protocol provided by the kit.

Plp1-CreER mice (Stock#: 005975) and *Rosa26*-LSL-tdTomato (Ai9/+) mice (Stock#: 007909) were from The Jackson Laboratory. The *Rosa26*-LSL-TAT/+ mouse strain was reported in our previous study (69). All mice were bred and raised in SPF-level animal rooms, and animal procedures were performed according to the guidelines (NA-032-2019) of the IACUC of the Institute of Neuroscience (ION), CAS Center for Excellence in Brain Science and Intelligence Technology, Chinese Academy of Sciences.

### Sample processing, histology and immunofluorescence assays, and cell counting

Mice were anesthetized and sacrificed, during which the heart was perfused with fresh 1× PBS and then with 4% paraformaldehyde (PFA) to remove blood from the inner ear. Subsequently, inner ear tissues were dissected out, post-fixed with 4% PFA overnight at 4°C, and washed thrice with 1× PBS, after which the inner ear samples were decalcified with 120 mM EDTA (Cat#: ST066, Beyotime) for 2 days at 4°C (until they were adequately soft) before whole-mount preparation. For whole-mount analysis, each cochlea was divided into three parts and first scanned at 10× magnification under a confocal microscope (Nikon C2, TiE-A1, or NiE-A1 Plus). In each acquired image, a line was drawn passing through the center of the IHCs and OHCs to measure the entire length of each cochlear duct, and the cochlear sample was then precisely divided into basal, middle, and apical portions of equal length.

The following primary antibodies were used for immunolabeling: anti-Prestin (goat, 1:1,000, sc-22692, Santa Cruz), anti-vGlut3 (rabbit, 1:500, 135203, Synaptic Systems), anti-Myo7a (rabbit, 1:500, 25-6790, Proteus Bioscience), anti-Ctbp2 (mouse, 1:200, 612044, BD Biosciences), anti-Fabp7 (rabbit, 1:500, ab32423, Abcam), and anti-NF- 200 (mouse, 1:500, N0142, Sigma). Cochlear tissues were counterstained with Hoechst 33342 solution (1:1,000, H3570, Thermo Scientific) to visualize nuclei. Lastly, cochlear samples were mounted with Prolong Gold antifade medium (P36930, Thermo Scientific). Immunofluorescence labeling was performed as described in our previous protocol (70).

### Quantification of endogenous and new IHCs, cell-fate conversion rate, and ribbon synapses

First, Myo7a+ cells were quantified in cochleae of control (no DT) and DT-treated *Fgf8-DTR/+* mice at P42 (Figure 1) by scanning entire cochlear turns under a confocal microscope at 10× magnification. Second, numbers of new IHCs (tdTomato+/vGlut3+) or tdTomato+ cells were estimated through confocal scanning at 60× magnification with 1 μm interval. For each turn, three areas were selected and a final average cell number was obtained per sample, after which a second average was calculated among all mice analyzed. Cell-fate conversion rate was calculated by normalizing tdTomato+/vGlut3+ cell numbers against total tdTomato+ cell numbers in the same confocal scanning regions. Notably, we only quantified tdTomato+ cells that were near vGlut3+ IHCs, which allowed us to confidently define these cells as derivatives of IBCs/IPhs regardless of their cell fates.

To precisely quantify Ctbp2+ ribbon synapses, three frequency areas (8, 16, and 30 kHz) were selected according to a previous method (71). Confocal scanning with 2.0× digital zoom and 0.41 μm interval was performed on samples from WT or Plp1-TAT- DTR mice at P42 (Figure 2E-F’). Ctbp2+ puncta were manually counted, and the Myo7a or tdTomato signal helped mark the boundary of each IHC. All cell-counting data are presented as mean ± SEM, and statistical analyses were performed using one-way ANOVA, followed by a Student’s *t* test with Bonferroni correction.

### Preparation of single-cell suspensions for single-cell RNA-seq

For 10× Genomics single-cell RNA-seq, fresh cochleae of Plp1-Ai9 mice (for P42_IBCs/IPhs) and Plp1-TAT-DTR mice (for P42_new IHCs) were dissected out and first digested in a choline chloride solution containing 20 U/mL papain (Cat#: LK003178, Worthington) and 100 U/mL DNase I (Cat#: LK003172, Worthington) for 20 min at 37°C, and then digested again with an additional protease (Cat#: P5147, Sigma; 1 mg/mL) and dispase (Cat#: LS02104, Worthington; 1 mg/mL) for 20 min at 25°C. The choline chloride solution contained 92 mM choline chloride, 2.5 mM KCl, 1.2 mM NaH_2_PO_4_, 30 mM NaHCO_3_, 20 mM HEPES, 25 mM glucose, 5 mM sodium ascorbate, 2 mM thiourea, 3 mM sodium pyruvate, 10 mM MgSO_4_.7H_2_O, 0.5 mM CaCl_2_.2H_2_O, and 12 mM N-acetyl-L-cysteine. Next, the post-digested tissues were gently triturated using fire-polished glass Pasteur pipettes (Cat#: 13-678-20b, Fisher) featuring four distinct pore sizes (largest size used first, smallest last). Lastly, the obtained single-cell suspensions were filtered using a 30 μm cell strainer (Cat#: 130- 098-458, Miltenyi), and any cell debris was removed using a debris-removing buffer (Cat#: 130-109-398, Miltenyi). All procedures were completed within 2.5 h after the animals were euthanized. The quality and survival rate of single cells were estimated using trypan blue solution (Cat#: 15250061, Thermo Fisher Scientific) before the cells were subject to 10× Genomics single-cell preparation and library procedures (Chromium Single Cell 3’ v3), and paired-end sequencing was performed on the final libraries on an Illumina Novaseq platform; the average sequencing depth of Plp1-Ai9 and Plp1-TAT-DTR libraries was 111,228 and 84,165 reads, respectively. The 10× Genomics single-cell RNA-seq raw data *de novo* generated in this study are accessible in the GEO (Gene Expression Omnibus) database under Accession No. GSE188259.

### Bioinformatics analysis

For single-cell RNA-seq data produced using the 10× Genomics platform, establishment of mouse reference genome and alignment were performed using Cell Ranger (v3.0.2) (72). Downstream analysis was performed using the R package Seurat (v3.2.3) (73). Cells with <500 or >7,500 unique genes and 20% mitochondrial genes were excluded from the analysis. The expression data were normalized using the “NormalizeData” function. Mitochondrial content and the effects of cell cycle heterogeneity were regressed out when scaling data in the “ScaleData” function. Principal component analysis was conducted using “RunPCA” and clustering was performed using “FindNeighbors” and “FindClusters.” Two types of dimensional reduction, uniform manifold approximation and projection (UMAP) and t-distributed stochastic neighbor embedding (t-SNE), were conducted using “RunUMAP” and “RunTSNE,” respectively. To integrate data from the 10× Genomics platform and smart-seq platform (from our previous two studies, GEO Accession Nos. GSE161156 and GSE199369), “FindIntegrationAnchors” and “IntegrateData” were applied for integration analysis.

P42_new IHCs were defined as cells in which the expression levels of *tdTomato* (WPRE sequence in Ai9 mouse strain), *Myo6*, *Myo7a*, and *Slc17a8* were above zero. Moreover, *Slc26a5* and *Ikzf2* were examined and their expression levels were found to be nearly zero in P42_new IHCs. *Slc1a3*+/*tdTomato*+/*Celf4*-/*Otof*-/*Fgfr3*- cells were defined as P42_IBCs/IPhs and *Fabp7*+/*Matn4*+/*Prox1*-/*Npy*- cells as P1_IBCs/IPhs. We also selected E16/P1/P7_IHCs from a previous study (GSE137299) (27). E16_IHCs and P7_IHCs were defined as *Myo6+/Fgf8+* cells, and *Myo6+/Fgf8+/Myo7a+* cells were classified as P1_IHCs.

To identify cell-type-specific genes, distinct criteria were set for cells sequenced using the 10× Genomics platform and the smart-seq platform. Differentially expressed genes (DEGs) among P42_IBCs/IPhs and other cell types were regarded as the initial P42_IBC/IPh genes (436 genes in Supplemental Figure 5D), which were calculated by employing “FindAllMarkers” with a log_2_ fold-change (FC) of ≥0.5 and adjusted p-value of ≤0.05 by using the Wilcoxon rank-sum test. P30_IHC and P60_PC/DC genes were initially selected with an average TPM of >16, and at least half the cells expressed these genes with a TPM value of >16. To further narrow down the P30_IHC genes, we overlapped our P30_IHC genes (single-cell smart-seq data of GSE199369) with the P30_IHC genes of a bulk-seq database (GSE111348) from a previous study (50); the shared 2,226 genes are listed in Supplemental Figure 5B. After subtracting the 1,781 overlapping genes (Supplemental Figure 5C) between P30_IHC and P60_PC/DC genes, 445 P30_IHC genes remained. After further discarding 13 genes that overlapped between the 445 P30_IHC and 436 P42_IBC/IPh genes, we identified 432 P30_IHC genes and 423 P42_ IBC/IPh genes. For P42_new IHCs, a gene was regarded as being highly expressed if (1) the gene’s average normalized expression value was >0.5, and (2) >50% of the cells expressed the gene with a normalized expression value of >0.5.

DEGs between P42_new IHCs and P42_IBC/IPhs were calculated using “FindMarkers” (log_2_FC>0.5, p<0.05). Based on the identified DEGs, gene ontology (GO) terms of biological process enrichment were evaluated (FDR<0.05) using DAVID (Database for Annotation, Visualization, and Integrated Discovery). By using the pre- processed matrix in the Seurat object, trajectory analysis was performed via Monocle (v2.14.0) (74). The top 2,000 most variable genes calculated using the “FindVariableFeatures” function in Seurat were used to construct a pseudotime trajectory, and the “orderCells” function in Monocle was used to arrange cells along the pseudotime axis.

### ABR measurement

ABR values were measured at 4, 5.6, 8, 11.3, 16, 22.6, 32, and 45 kHz at P42, following our previously published protocol (10). Student’s *t* test was applied to determine the statistical significance regarding the hearing thresholds at the same frequency among different mice (Figures 1E and 3F).

### SEM and TEM sample preparation and analysis

We followed the SEM protocol detailed in our previous study (34). In TEM analysis of IHCs, similar sample-treatment steps were used as those for SEM, except that the samples were subsequently embedded in epoxy resin monomer (Cat#: 90529- 77-4, SPI Supplies); lastly, 70 nm ultrathin transverse sections of the OC were obtained using an ultramicrotome (Leica EM UC6, Germany) and examined using a JEOL-1230 transmission electron microscope (Nippon Tekno, Japan).

### Calcium current and exocytosis measurement in WT and new IHCs

The apical turns of cochlear sensory regions were dissected and then pinned on a glass coverslip. All recordings were performed at room temperature. Recording pipettes were pulled from borosilicate glass capillaries (Cat#: TW150-4F, World Precision Instruments) and coated with dental wax. The typical pipette-resistance range was 4–7 MΩ. IHCs were voltage-clamped using an Axon 200B amplifier (Axon Instruments- Molecular Devices) interfaced with a Digidata 1440B (Molecular Devices, LLC.). All recordings were performed by using the jClamp software (details please refer to http://www.scisoftco.com/jclamp.html, New Haven, CT, USA). Liquid junction potential was corrected offline.

For recording Ca^2+^ currents and exocytosis, the extracellular solution contained (in mM) 105 NaCl, 2.8 KCl, 30 TEA-Cl, 5 CaCl_2_, 1 MgCl_2_, 2 Na-pyruvate, 1 creatine, 10 D-glucose, and 10 HEPES (pH 7.4 with NaOH), with D-glucose used to adjust osmolarity to 300 mOsm; and the pipette solution contained (in mM) 105 Cs-methane sulfonate, 20 CsCl, 10 HEPES, 10 TEA-Cl, 1 EGTA, 4 Mg-ATP, 0.5 Na-GTP, and 5 phosphocreatine-Na (pH 7.2 with CsOH), with D-glucose used to adjust osmolarity to 298–300 mOsm. A 500 ms voltage ramp from -80 to +70 mV was delivered to IHCs to record the Ca^2+^ current (ICa). To analyze calcium channel activation, conductance-voltage relationships were calculated from the Ca^2+^ current responses to ramp stimulation and fitted with the Boltzmann equation:

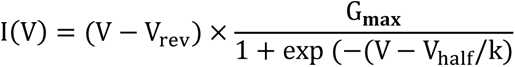

Here, V is the command membrane potential, G_max_ is the maximum conductance, V_half_ is the half-activation voltage, and k is the slope factor that defines the steepness of voltage dependence in current activation.

Membrane capacitance changes were recorded to monitor the fusion of synaptic vesicles during exocytosis. IHCs were held at -80 mV, and exocytosis was induced by applying a depolarizing pulse of 0 mV. Sinewaves of 1 kHz and 70 mV peak-to-peak amplitude were superimposed on the holding potential before and after stimulation.

### MET current measurement in WT and new IHCs

Whole-cell patch-clamp was used to record the MET current of IHCs, according to protocols described previously (75). Adult cochlear sensory regions were dissected in a solution containing (in mM) 141.7 NaCl, 5.36 KCl, 0.1 CaCl_2_, 1 MgCl_2_, 0.5 MgSO_4_, 3.4 L-glutamine, 10 glucose, and 10 H-HEPES (pH 7.4), and then transferred into a clean recording chamber bathed with a recording solution containing (in mM) 144 NaCl, 0.7 NaH_2_PO_4_, 5.8 KCl, 1.3 CaCl_2_, 0.9 MgCl_2_, 5.6 glucose, and 10 H-HEPES (pH 7.4).

Patch pipettes featuring a tip diameter of 2–3 μm were prepared using a pipette puller (Sutter, P-1000). The intracellular solution contained (in mM) 140 KCl, 1 MgCl_2_, 0.1 EGTA, 2 Mg-ATP, 0.3 Na-GTP, and 10 H-HEPES (pH 7.2). The MET currents of IHCs were sampled at 100 kHz with a patch-clamp amplifier (Axon, 700B), and the holding potential was set at -70 mV. To evoke MET currents, a fluidjet pipette featuring a tip diameter of 6–10 μm was positioned near a hair bundle (at a distance of ∼5 μm), and 40 Hz sine waves were delivered through a piezoelectric disc to stimulate IHC hair bundles.

## ACKNOWLEDGMENTS

We thank Dr. Min Zhang and Ms. Zhenning Zhou from the Molecular Biology Facility of the Institute of Neuroscience (ION) for 10× Genomics single-cell library preparation; Drs. Xu Wang and Yu Kong from the Electronic Microscope Facility of the ION for SEM and TEM assistance; Dr. Qian Hu from the Optical Imaging Facility of the ION for support with image analysis; and Ms. Qian Liu from the Department of Embryology of the ION animal center for helping us in transplanting zygotes into pseudopregnant female mice. This study was funded by the National Key R&D Program of China (2021YFA1101804), National Natural Science Foundation of China (82000985 and 82101217), Strategic Priority Research Program of Chinese Academy of Science (XDB32060100), Shanghai Municipal Science and Technology Major Project (2018SHZDZX05), and Innovative Research Team of High-Level Local Universities in Shanghai (SSMU-ZLCX20180601).

## SUPPLEMENTAL FIGURE LEGENDS

**Supplemental Figure 1.**
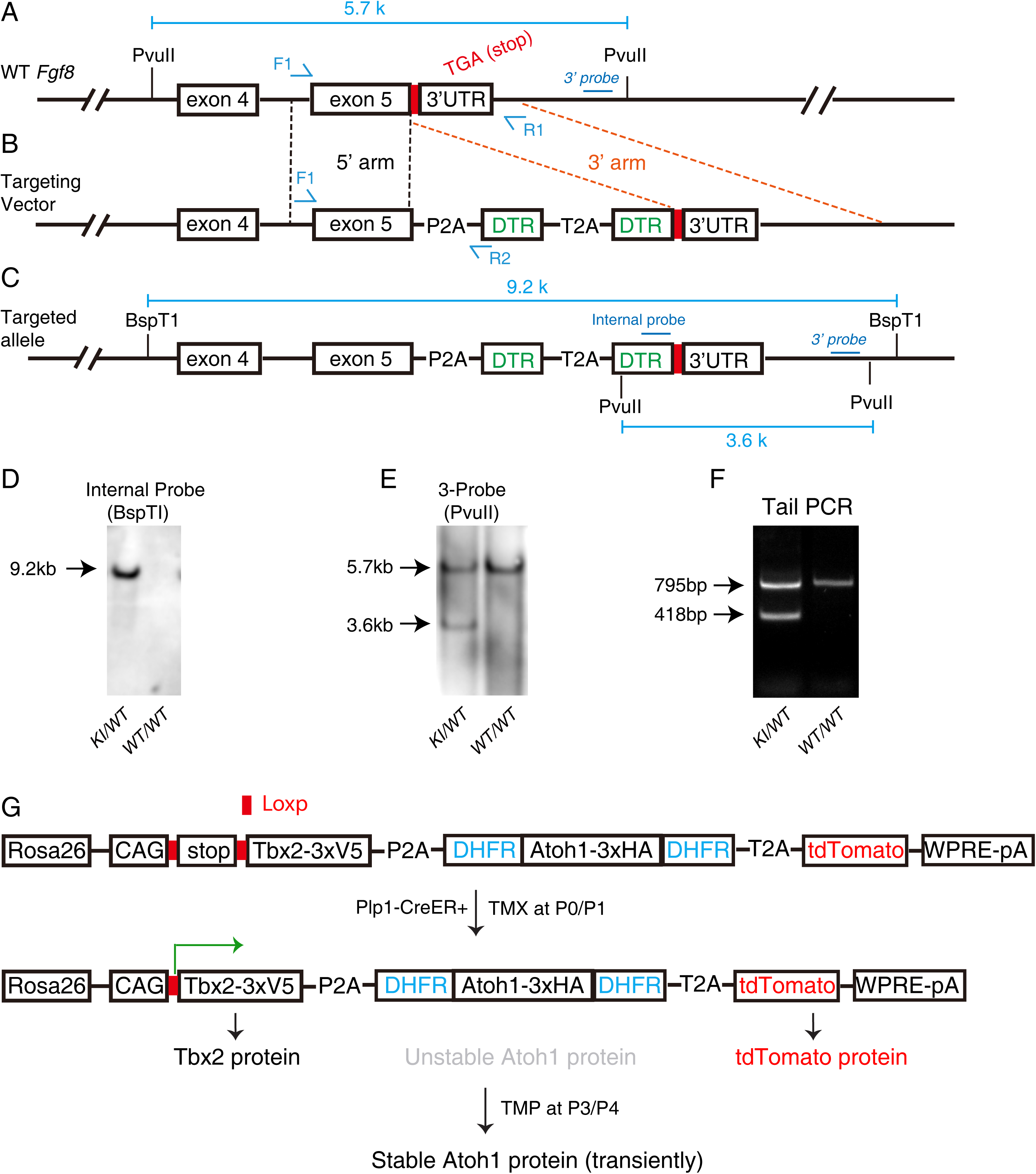
Generation of new *Fgf8*-DTR/+ mice. **(A-C)** Immediately before stop-codon TGA (red in A) in wild-type (WT) *Fgf8*, we inserted an element containing P2A-DTR-T2A-DTR (B); final post-targeted *Fgf8* allele is illustrated in (C). **(D-E)** Southern blotting with internal (D) and external 3ʹ (E) probes to confirm absence of random insertion of targeting vector in *Fgf8*-DTR/+ genome. **(F)** Tail-DNA PCR readily distinguished WT (795 bp) and knockin (KI; 418 bp) alleles. **(G)** Simple illustration of how Tbx2 and tdTomato proteins are permanently expressed after TMX induction and how transient Atoh1 protein expression is additionally induced when TMP is administered.

**Supplemental Figure 2.**
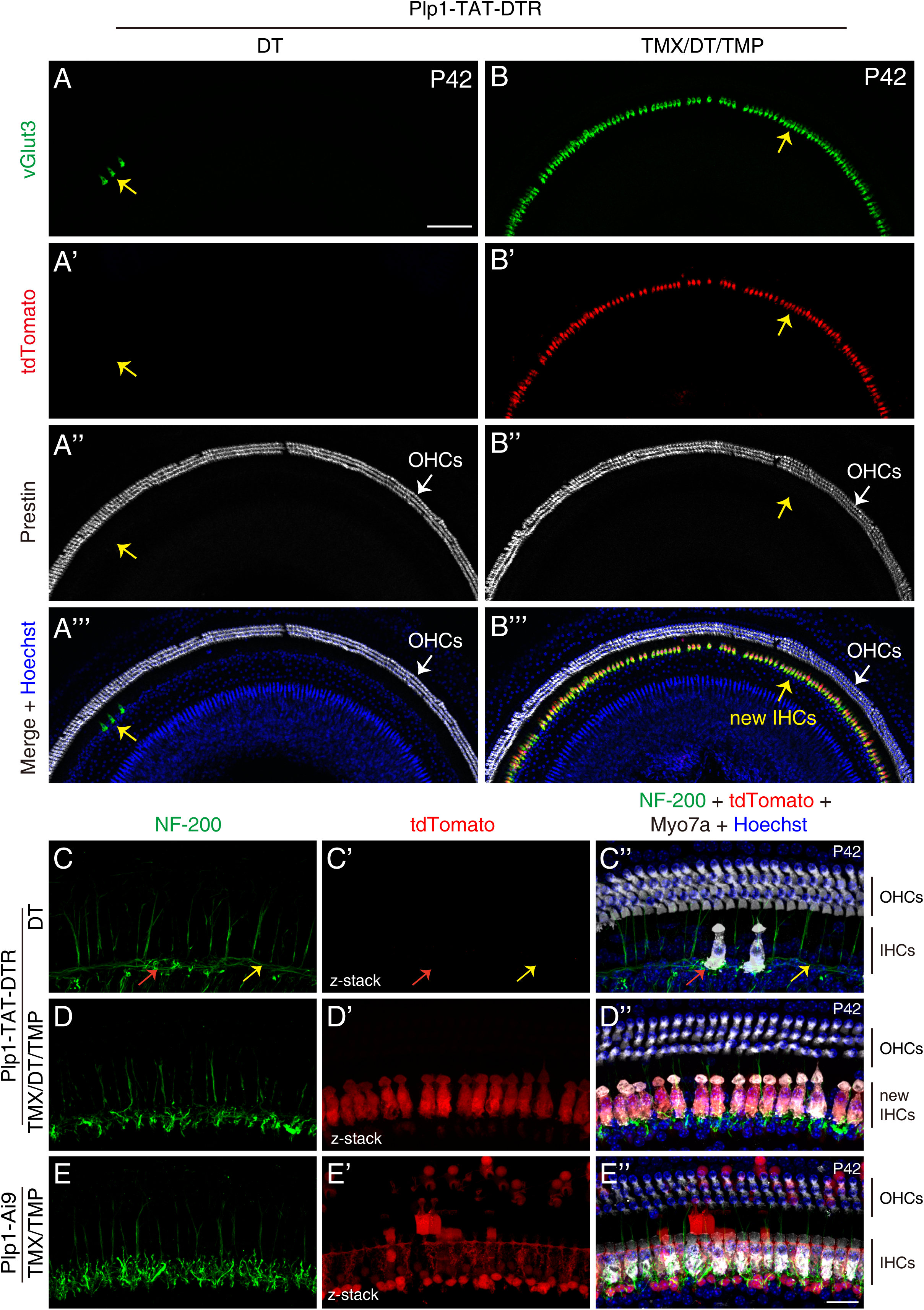
vGlut3+ new IHCs are innervated by SGN neural fibers. **(A-B’’’)** Triple labeling for vGlut3, tdTomato, and Prestin in cochleae of Plp1-TAT- DTR mice treated with DT only (A-A’’’) or TMX, DT, and TMP (B-B’’’). Arrows in (A-A’’’): endogenous IHCs remaining after DT treatment; arrows in (B-B’’’): one row of new IHCs that were tdTomato+/vGlut3+/Prestin-. Corresponding high-magnification images are presented in Figure 2B-C’’. **(C-E’’)** Triple labeling for NF-200, tdTomato, and Myo7a in cochleae of Plp1-TAT-DTR mice (C-D’’) and control Plp1-Ai9 mice (E-E’’). (C-C’’) Red arrows: one endogenous IHC remaining in Plp1- TAT-DTR mice treated with DT only; yellow arrows: region where IHCs were lost. Scale bars: 200 μm (A), 20 μm (E’’).

**Supplemental Figure 3.**
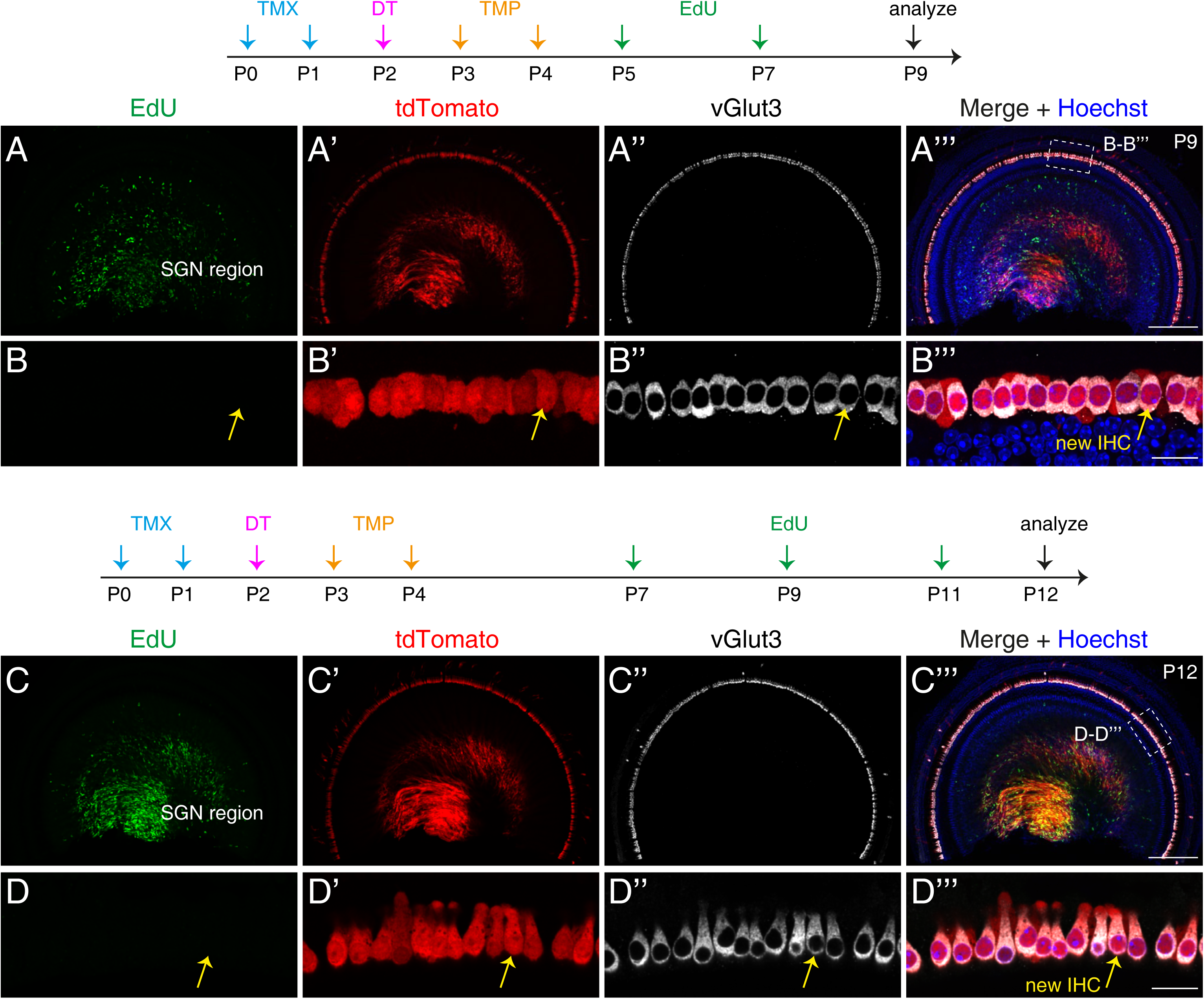
New IHCs are produced by direct transdifferentiation. Plp1-TAT-DTR mice were divided into two groups and subject to two experimental paradigms, as illustrated in (A-B’’’) and (C-D’’’). Cochlear samples were triple labeled with EdU, tdTomato, and anti-vGlut3. Dotted boxes in (A’’’) and (C’’’): areas shown at high resolution in (B-B’’’) and (D-D’’’). Yellow arrows: new IHCs positive for tdTomato and vGlut3 but not EdU at P9 (B-B’’’) and P12 (D-D’’’). Scale bars: 200 μm (A’’’ and C’’’), 20 μm (B’’’ and D’’’).

**Supplemental Figure 4.**
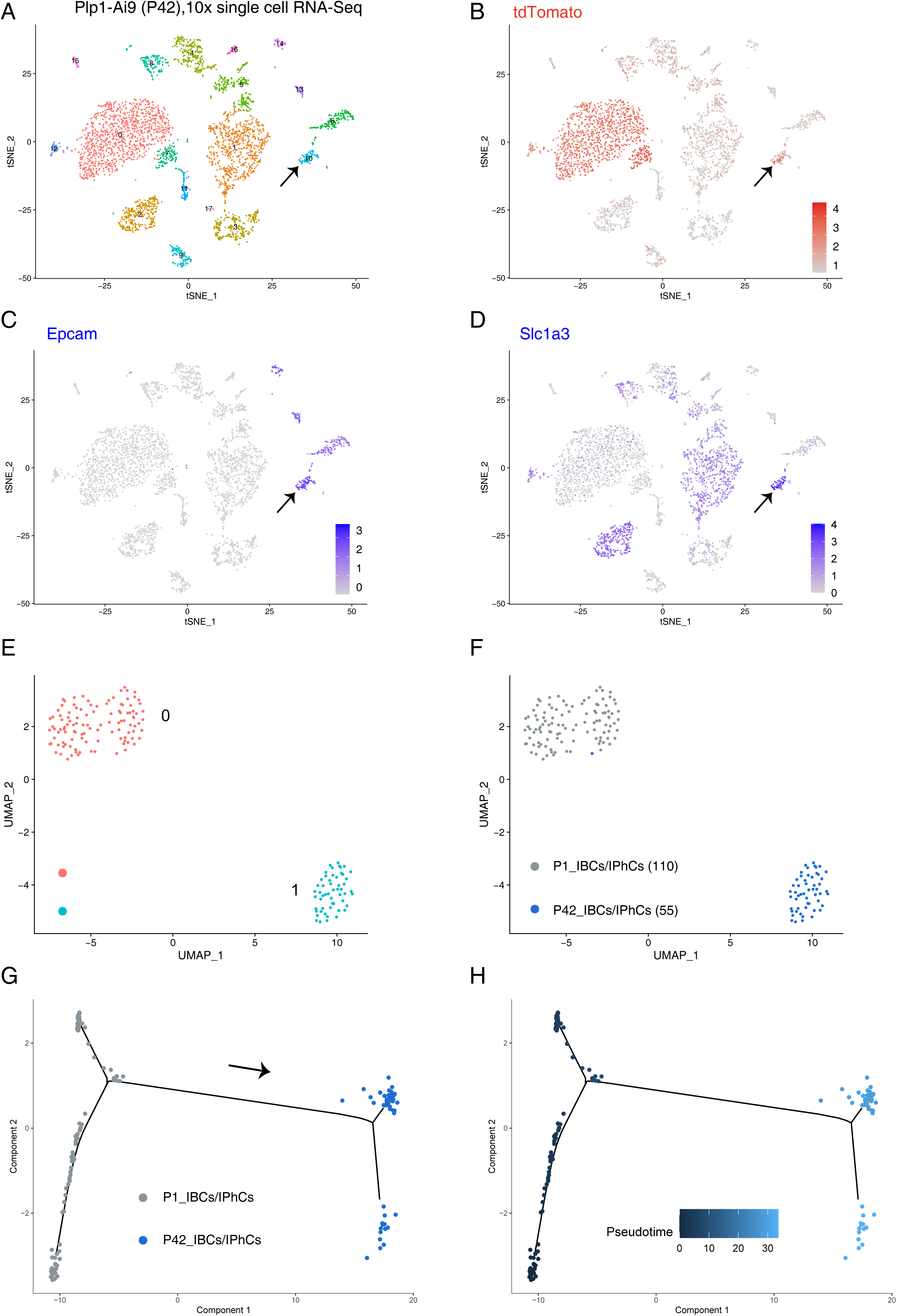
Single-cell RNA-seq of IBCs/IPhs. **(A-D)** Cochleae of Plp1- Ai9 mice at P42 were subject to 10× Genomics single-cell RNA-seq, and 18 cell clusters were identified (A). Arrows in (A-D): Cluster 10 containing 55 P42_IBCs/IPhs showing enrichment of *tdTomato* (B), *Epcam* (C), and *Slc1a3* (D). **(E-F)** UMAP analysis of 110 P1_IBCs/IPhs and 55 P42_IBCs/IPhs. Two clusters were formed (E), and cells at the same age were aggregated (F). **(G-H)** Trajectory analysis of P1_IBCs/IPhs and P42_IBCs/IPhs. Arrow in (G): developmental direction from P1 to P42. The calculated pseudotime matched the intrinsic ages (H).

**Supplemental Figure 5.**
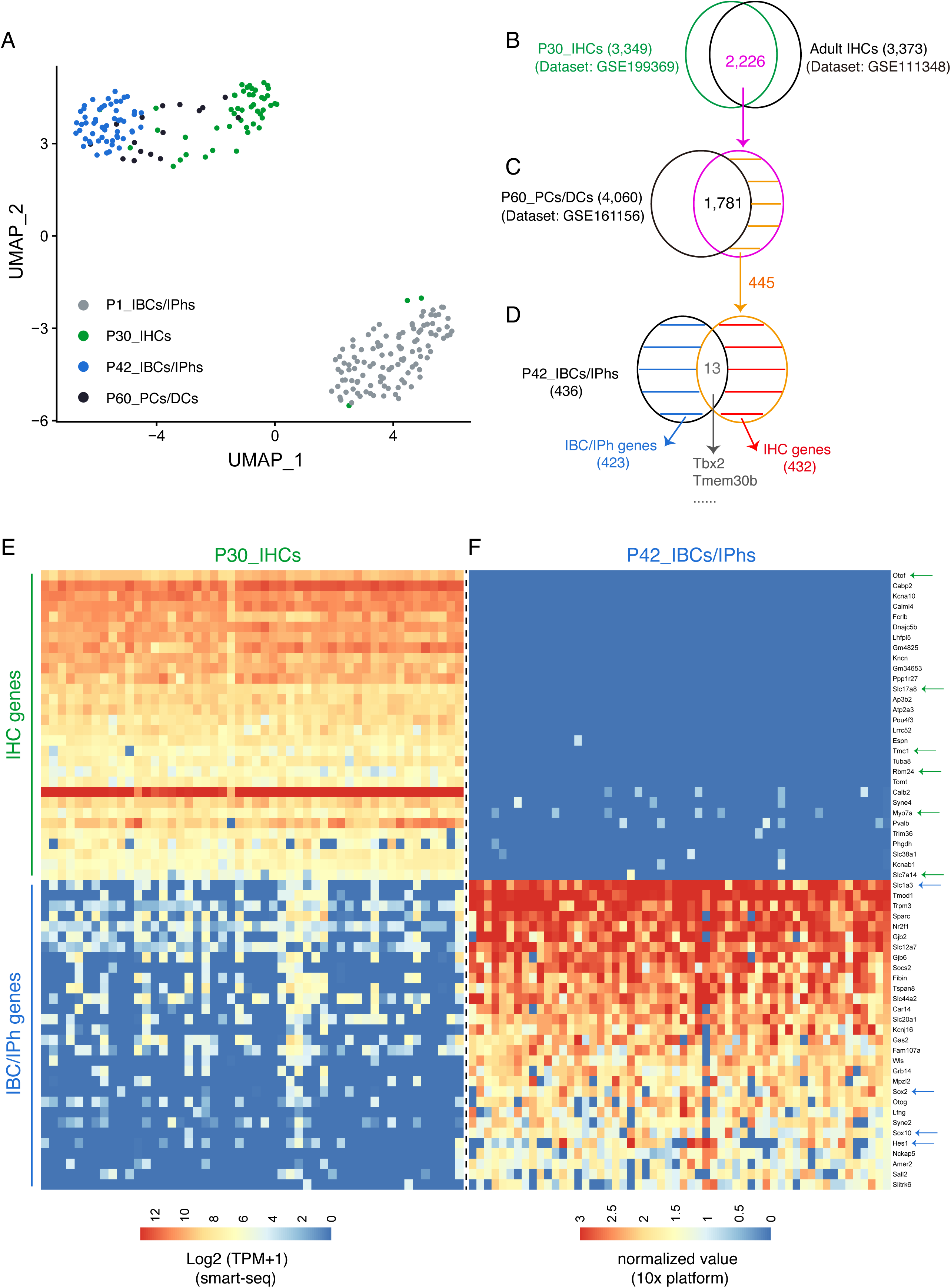
Identification of genes enriched in adult IHCs and IBCs/IPhs. **(A)** UMAP analysis of P1_IBCs/IPhs, P42_IBCs/IPhs, P30_IHCs, and P60_PCs/DCs. Majority of P30_IHCs clustered together. **(B-D)** Illustration of our step-by-step approach to define the final 432 IHC and 423 IBC/IPh genes; 2,226 genes were detected at a high expression level in both P30_IHCs here (green circle in B) and adult IHCs from another study (black circle in B). Among the 2,226 genes, 1,781 were also highly expressed in P60_PCs/DCs from our previous study (black circle in C), and 13 out of the remaining 445 genes were also included in the 436 genes enriched in P42_IBCs/IPhs (black circle in D). **(E-F)** Plotting of top IHC and IBC/IPh genes in P30_IHCs (E) and P42_IBCs/IPhs (F). Each row shares the same gene listed on the right side, but (E) and (F) were independently processed using distinct bioinformatics approaches and the gene-expression levels are presented with different scales (bottom). IHC genes (green arrows) included IHC-specific genes such as *Otof*, *Slc17a8*, and *Slc7a14*, as well as pan-HC genes such as *Tmc1*, *Rbm24*, and *Myo7a*. IBC/IPh genes included *Slc1a3*, a known IBC/IPh gene, and pan-SC genes such as *Sox2* and *Sox10* (blue arrows).

**Supplemental Figure 6.**
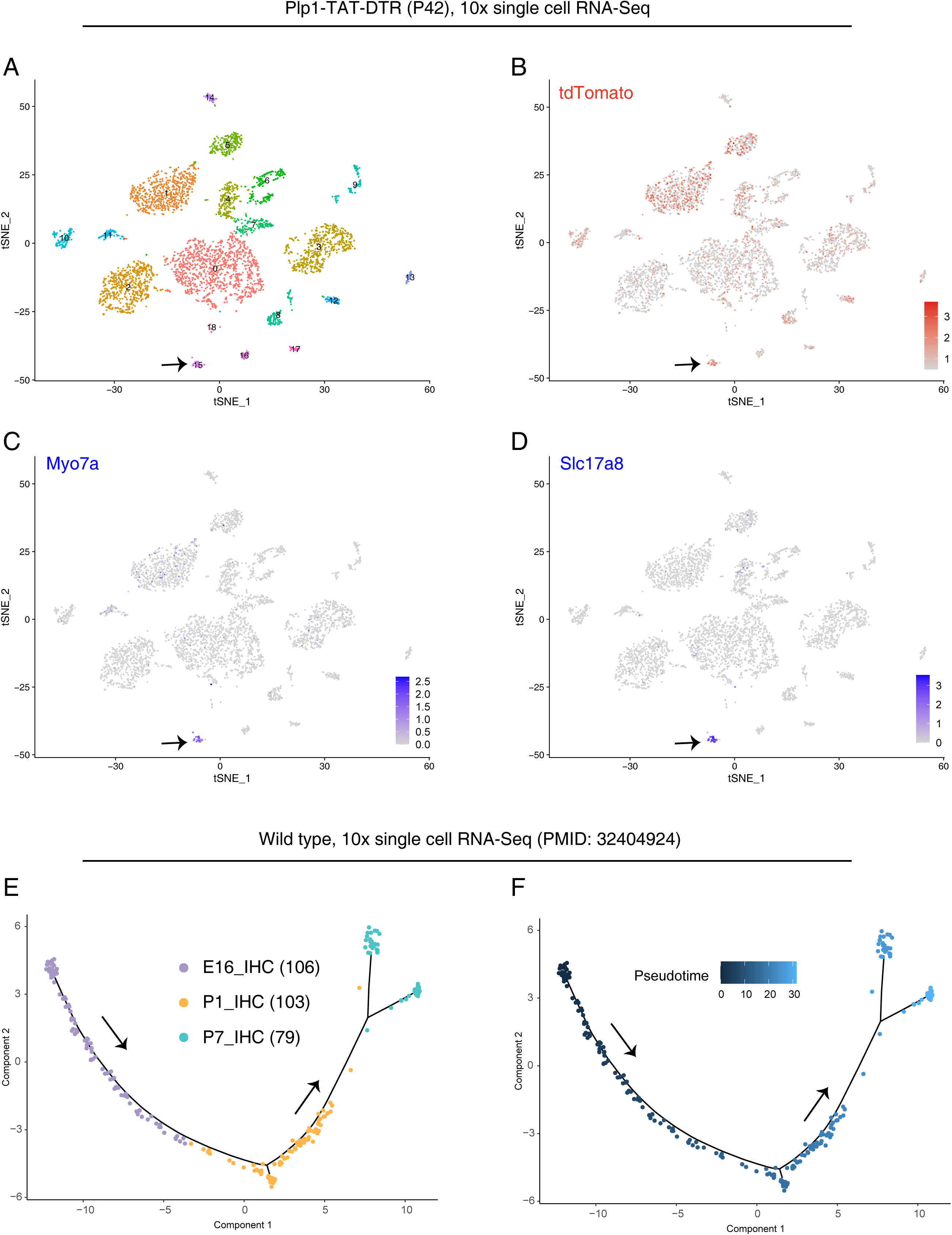
Single-cell RNA-seq of new IHCs. **(A-D)** Cochleae of Plp1- TAT-DTR mice treated with TMX, DT, and TMP were dissected at P42 and subject to 10× Genomics single-cell RNA-seq; 19 cell clusters were identified (A). Arrows in (A-D): Cluster 15 containing the 42 new IHCs identified. Notably, *tdTomato* (B), *Myo7a* (C), and *Slc17a8* (D) were enriched in the P42_new IHCs. **(E-F)** Trajectory analysis of endogenous IHCs at E16, P1, and P7 by using data from one previous study. Arrows: developmental direction from E16 to P7 (E). The calculated pseudotime matched the intrinsic time (F).

**Supplemental Figure 7.**
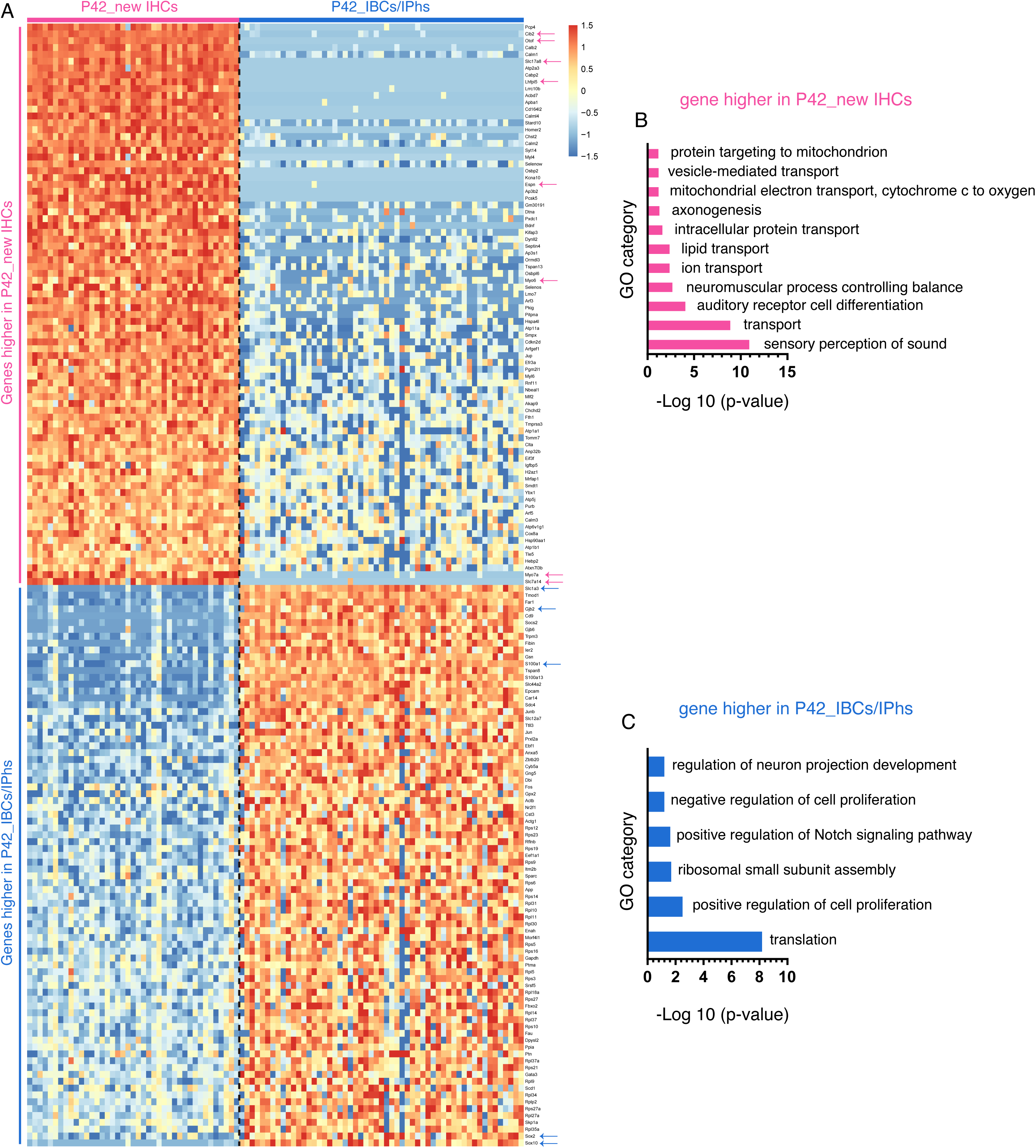
New IHCs upregulate 183 IHC genes and downregulate 191 IBC/IPh genes. **(A)** Plot of top differentially expressed genes between new P42_IHCs and P42_IBCs/IPhs. Top half: genes expressed at significantly higher levels in P42_new IHCs than in P42_IBCs/IPhs. Pink arrows: IHC-specific genes *Otof*, *Slc17a8*, and *Slc7a14* and pan-HC genes *Cib2*, *Lhfpl5*, *Espn*, *Myo6*, *Lmo7*, and *Myo7a*. Bottom half: genes expressed at markedly lower levels in P42_new IHCs than in P42_IBCs/IPhs. Blue arrows: IBC/IPh genes such as *Slc1a3*, *Gjb2*, *S100a1*, *Sox2*, and *Sox10*. **(B-C)** GO analysis of genes expressed at higher (B) and lower (C) levels in P42_new IHCs than in P42_IBCs/IPhs.

**Supplemental Figure 8.**
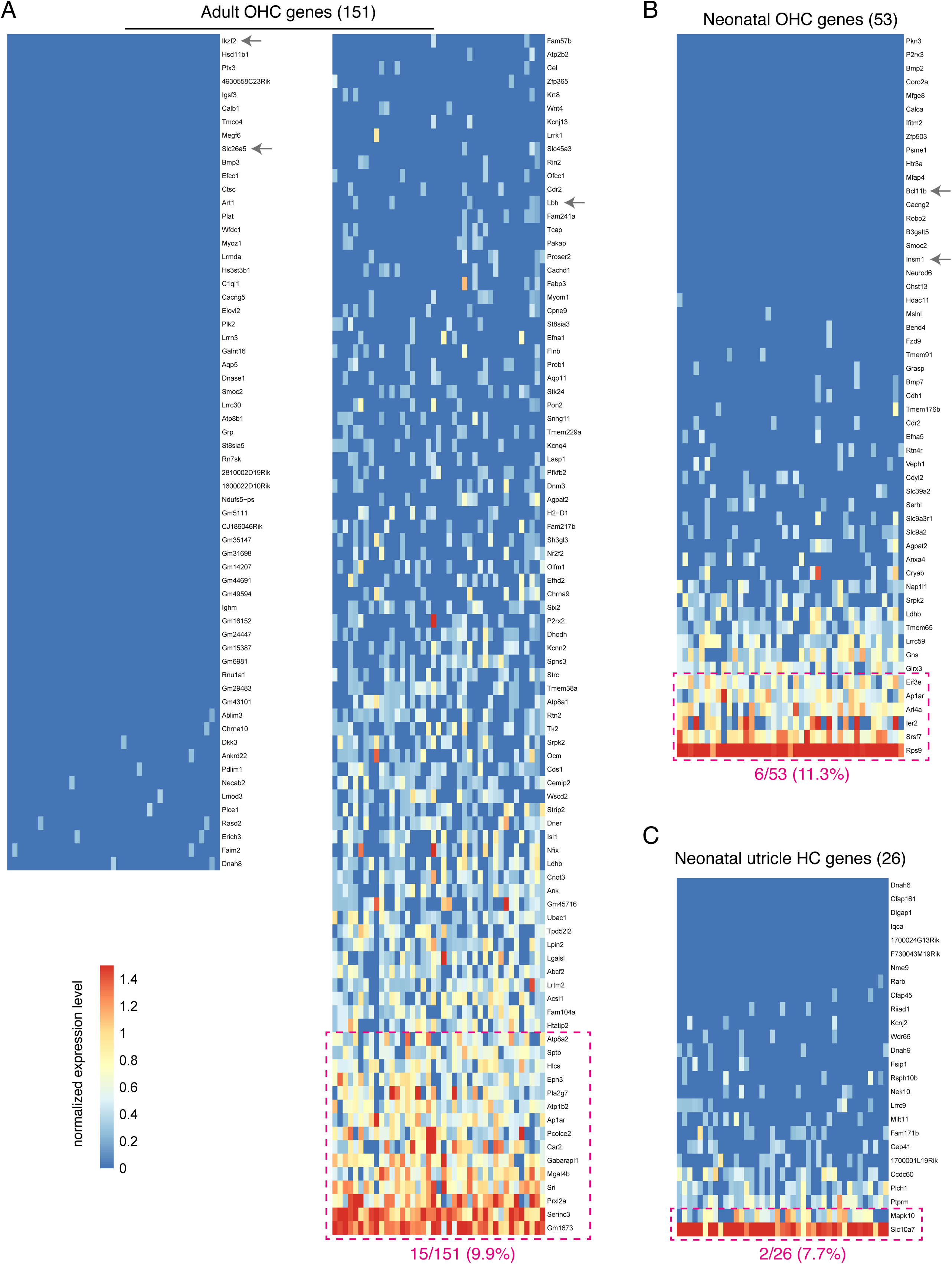
OHC and utricle genes are minimally expressed in new IHCs. **(A)** Plot of 166 adult OHC genes, including *Ikzf2*, *Slc26a5*, and *Lbh* (gray arrows); dotted pink box: 15/151 (9.9%) genes regarded as being expressed in new IHCs. **(B)** Plot of 53 neonatal OHC genes, including *Bcl11b* and *Insm1* (gray arrows); dotted pink box: 6/53 (11.3%) genes defined as being expressed in new IHCs. **(C)** Plot of 26 neonatal utricle HC genes; dotted pink box: only 2/26 (7.7%) genes were expressed in new IHCs.

**Supplemental Figure 9.**
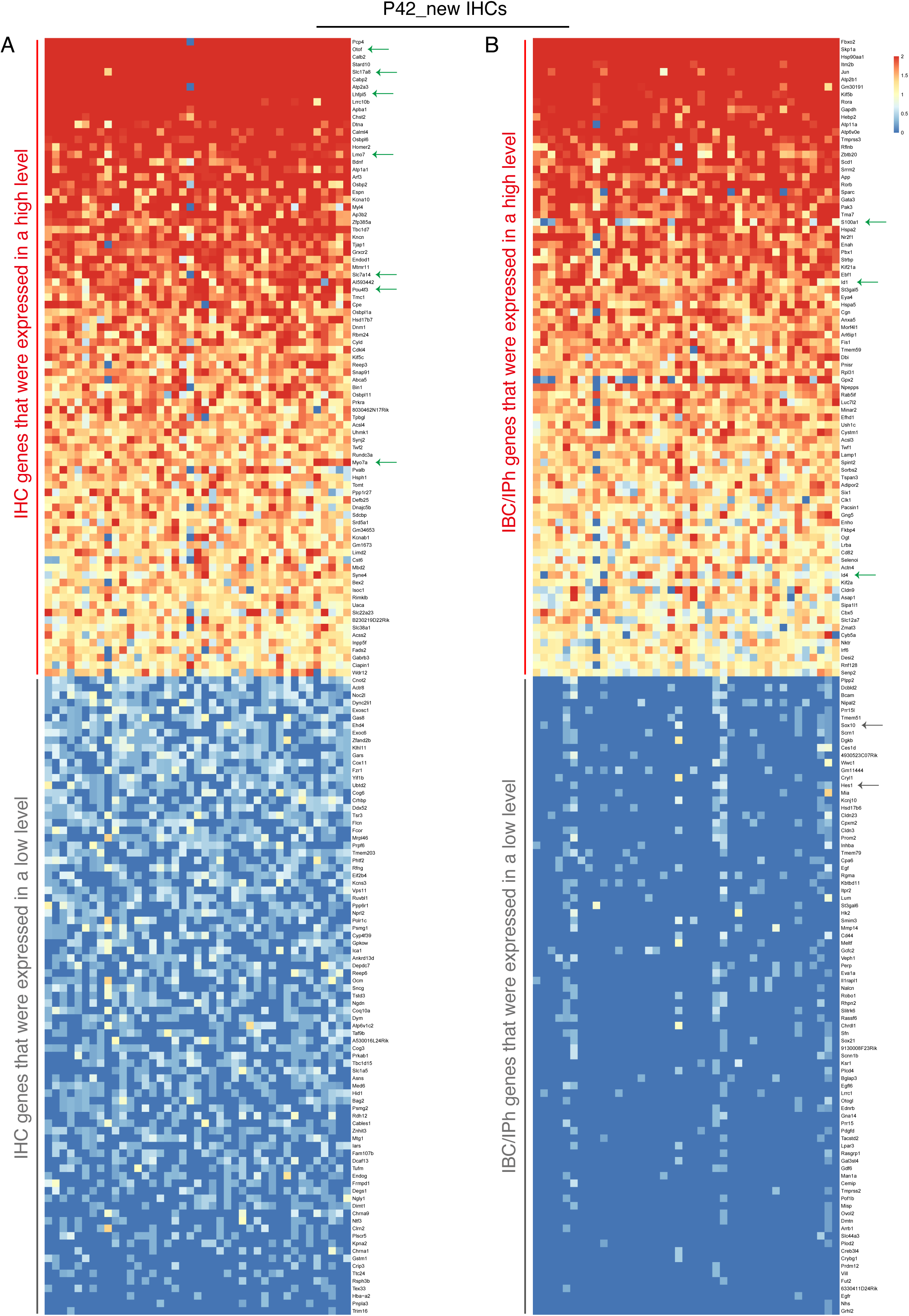
Approximately 40% of IHC and IBC/IPh genes are not upregulated and downregulated, respectively, in new IHCs. **(A)** Sampled IHC genes expressed at high (red line) or low (gray line) levels. Green arrows: a few well-known IHC-specific genes (e.g., *Slc17a8* and *Otof*) or pan-HC genes (e.g., *Myo7a* and *Pou4f3*). **(B)** Sampled IBC/IPh genes expressed at high (red line) or low (gray line) levels. Green arrows: a few well-known IBC/IPh genes (e.g., *S100a1*, *Id1*, and *Id4*) expressed at high levels; gray arrows: IBC/IPh genes (*Sox10* and *Hes1*) expressed at low levels.

